# An atypical E3 ligase safeguards the ribosome during nutrient stress

**DOI:** 10.1101/2024.10.10.617692

**Authors:** Todd Douglas, Jiasheng Zhang, Zhiping Wu, Kyrillos Abdallah, Meghan McReynolds, Wendy V. Gilbert, Kazuhiro Iwai, Junmin Peng, Lawrence H. Young, Craig M. Crews

**Affiliations:** Department of Molecular, Cellular, Developmental Biology, Yale University, New Haven, CT, USA; Yale Cardiovascular Research Center, Yale School of Medicine, New Haven, CT, USA; Departments of Structural Biology and Developmental Neurobiology, St. Jude Children’s Research Hospital, Memphis, TN, USA; Department of Molecular Biophysics and Biochemistry, Yale University, New Haven, CT, USA; Department of Molecular and Cellular Physiology, Graduate School of Medicine, Kyoto University, Kyoto, Japan; Department of Molecular and Cellular Physiology, Yale University, New Haven, CT, USA; Department of Pharmacology, Yale University, New Haven, CT, USA; Department of Chemistry, Yale University, New Haven, CT, USA

**Keywords:** HOIL-1, metabolic stress, ribotoxic stress, cell death, ubiquitin, signaling, cardiomyopathy

## Abstract

Metabolic stress must be effectively mitigated for the survival of cells and organisms. Ribosomes have emerged as signaling hubs that sense metabolic perturbations and coordinate responses that either restore homeostasis or trigger cell death. As yet, the mechanisms governing these cell fate decisions are not well understood. Here, we report an unexpected role for the atypical E3 ligase HOIL-1 in safeguarding the ribosome. We find HOIL-1 mutations associated with cardiomyopathy broadly sensitize cells to nutrient and translational stress. These signals converge on the ribotoxic stress sentinel ZAKα. Mechanistically, mutant HOIL-1 excludes a ribosome quality control E3 ligase from its functional complex and remodels the ribosome ubiquitin landscape. This quality control failure renders glucose starvation ribotoxic, precipitating a ZAKα-ATF4-xCT-driven noncanonical cell death. We further show HOIL-1 loss exacerbates cardiac dysfunction under pressure overload. These data reveal an unrecognized ribosome signaling axis and a molecular circuit controlling cell fate during nutrient stress.

## INTRODUCTION

All living things must at some point mobilize their metabolic reserves to survive. Under conditions of scarcity, such as nutrient deprivation, cells reprogram their metabolism and become catabolic via the breakdown of stored fatty acids or glycogen. When these threatening states persist, cells engage stress response programs to restore homeostasis. Many such threats converge on the integrated stress response (ISR), during which global protein translation is suppressed while the transcription factor ATF4 is selectively translated^1^. ATF4 then orchestrates a transcriptional program to mitigate the stress. As the site of protein translation, ribosomes are now recognized as signaling hubs that sense a wide range of perturbations and coordinate appropriate effector responses^2–4^.

Failure of these programs to restore homeostasis results in the terminal stress response: cell death. Such cell fate decisions must be carefully controlled and employ failsafe mechanisms to prevent pathological cell loss, a key driver of a plethora of human diseases. The post-translational modification of proteins with ubiquitin plays a central role in coordinating cell fate. Accordingly, ubiquitin-related enzyme deficiency frequently manifests with excessive cell death and metabolic dysfunction^5^.

HOIL-1 is a RING-between-RING (RBR) family E3 ligase whose loss-of-function results in a fatal multi-organ disease with both immunological and metabolic symptoms^6–15^. This disease, termed polyglucosan body myopathy 1 with or without immunodeficiency (PGBM1), presents with a notable genotype-phenotype spectrum. Complete HOIL-1 loss yields fulminant autoinflammatory disease, while frameshift mutations that truncate the catalytic RBR domain predominantly cause myopathy^9^. The latter is associated with pronounced accumulation of glycogen aggregates in cardiomyocytes^7^, with heart failure secondary to dilated cardiomyopathy being the main cause of death.

HOIL-1 is most known for its role as a component of the linear ubiquitin assembly complex (LUBAC), an essential regulator of the immune response. LUBAC deficiency impairs immune cell activation and sensitizes to cell death *in vivo*^6,16–24^. Beyond LUBAC, HOIL-1 also has the rare ability to conjugate ubiquitin onto its substrates via an ester linkage. A recent landmark study demonstrated that HOIL-1 can directly ubiquitinate glycogen via this linkage *in vitro*^25^. In proteins, HOIL-1-mediated ester-linked ubiquitination occurs on serine and threonine residues and has been observed on several components of the MyD88 signalosome^26–28^. The immunological features of PGBM1 can be putatively ascribed to these established roles for HOIL-1 in immune signaling. On the other hand, the etiology of the fatal metabolic features of this disease is currently unknown.

While interrogating the role of HOIL-1 in glycogen metabolism, we herein discover that HOIL-1 plays a key role in safeguarding the ribosome during stress. We find that loss of the HOIL-1 RBR domain, as observed in a subset of PGBM1 patients, renders cells hypersensitive to glucose starvation-induced cell death. By characterizing this noncanonical cell death modality, we show it is initiated by the serine/threonine kinase ZAKα, a sensor of ribotoxic stress, and propagates via ATF4 through a mechanism distinct from that elicited by canonical ZAKα activators. We extend this to show HOIL-1^ΔRBR^ cells are broadly sensitive to ribotoxic stress. We define a molecular circuit wherein mutant HOIL-1 acts as a dominant negative that remodels the ribosome ubiquitin landscape, ultimately impairing a cytoprotective quality control response. These data position HOIL-1 as a previously unrecognized node in the ribosome signaling network.

## RESULTS

### HOIL-1 loss exacerbates cardiac remodeling and dilatation under pressure overload

Dilated cardiomyopathy is the most common feature of HOIL-1 loss-of-function in humans^15^. We first sought to establish whether this could be recapitulated in mice. Full HOIL-1 loss by targeting exons 1 and 2 of *Rbck1* (encoding HOIL-1) is embryonic lethal at e10.5 caused by unrestricted endothelial cell death^29^. Conversely, targeting exons 7 and 8 yields viable mice without overt disease, herein referred to as HOIL-1^null^ (ref. 30). Exons 7-12 encode the RBR domain, highlighting that deletion of the RBR produces a phenotype distinct from that of complete loss in mice and humans.

We subjected wildtype (WT; n=8) and HOIL-1^null^ (n=10) mice to transverse aortic constriction (TAC) surgery to induce pressure overload and assessed left ventricular (LV) hypertrophy and LV function and remodeling over 8 weeks (Figure 1A). HOIL-1^null^ mice presented with a normal baseline cardiac phenotype at 10 weeks of age. Despite similar pressure overload as indicated by aortic flow peak velocity after surgery (Figure 1B), HOIL-1^null^ males, but not females, developed more severe cardiac dilatation 4 weeks post-TAC (Figure 1C and 1D). Histology revealed an increase in cardiomyocyte cross-sectional area (male WT mean = 320.2 µm^2^, male HOIL-1^null^ mean = 409.5 µm^2^) and diameter (male WT mean = 17.87 µm, male HOIL-1^null^ mean = 20.37 µm), consistent with cellular hypertrophy (Figure 1E and 1F). As observed with PGBM1 patients, both male and female cardiomyocytes from HOIL-1^null^ mice accumulated dense glycogen aggregates that were absent from all WT mice examined (Figure 1G). We found no difference between genotypes in myocardial fibrosis using Masson’s trichrome staining of collagen (Figure 1H and 1I). LV contractile function, as assessed by LV ejection fraction, trended lower in male HOIL-1^null^ mice than WT mice but did not reach significance by 8 weeks (Figure 1J). Nevertheless, male HOIL-1^null^ mice had increased pulmonary edema as assessed by lung-to-body weight ratio (Figure 1K), indicative of elevated LV diastolic pressures from cardiac dysfunction. These data demonstrate that loss of HOIL-1 exacerbates cardiac remodeling with the development of excessive LV dilatation, cardiomyocyte hypertrophy, and heart failure following pressure overload in a sex-dependent manner.

**Figure 1.**
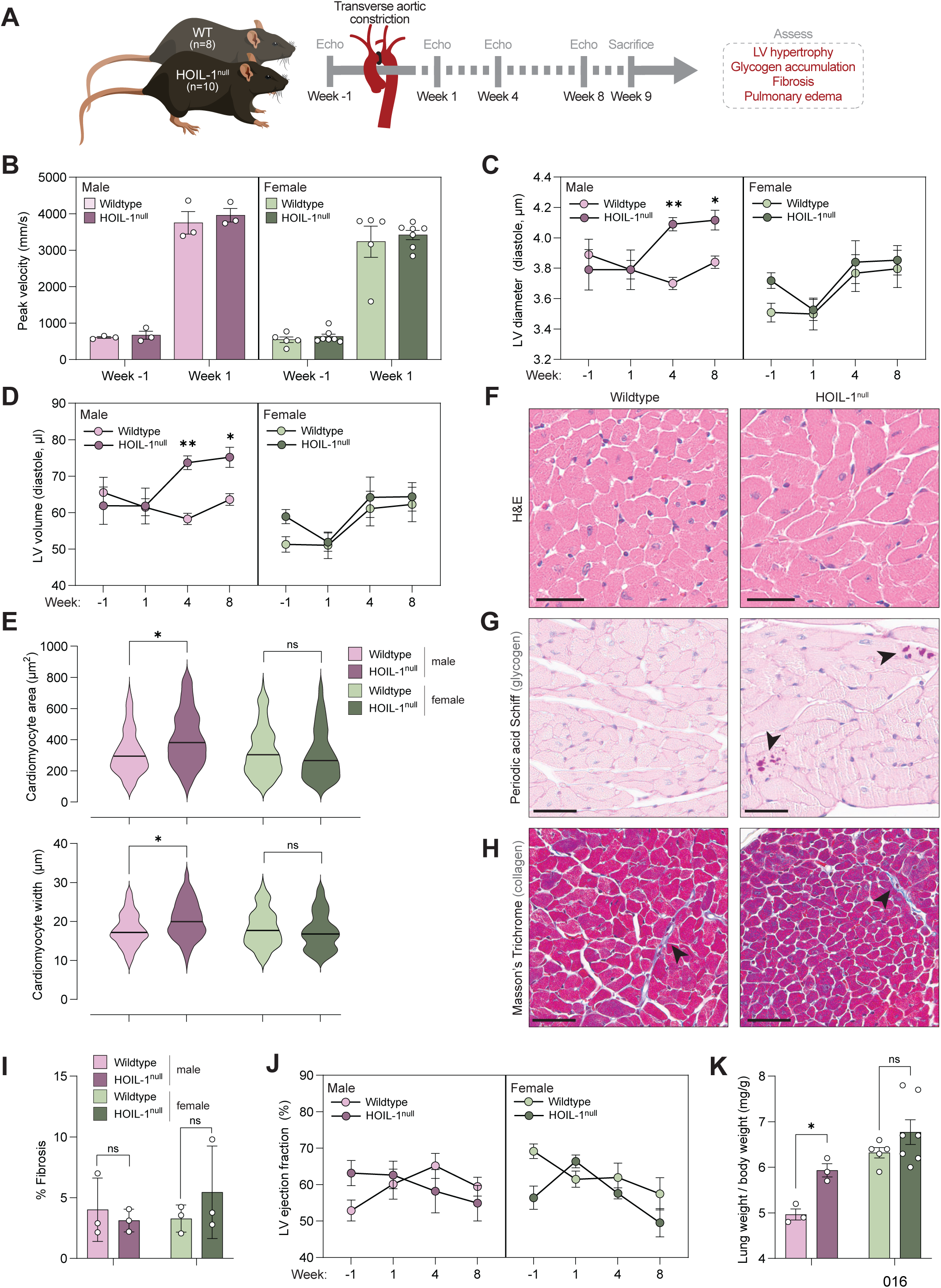
HOIL-1 loss exacerbates cardiac remodeling and hypertrophy under pressure overload. (A) Schematic of experimental design assessing cardiac hypertrophy in WT (n=8; 3 males, 5 females) and HOIL-1^null^ (n=10; 3 males, 7 females) mice following transverse aortic constriction (TAC) surgery. (B) Peak aortic flow velocity assessed by echocardiography 1 week pre- and post-TAC. (C) LV diameter assessed by echocardiography. (D) LV volume assessed by echocardiography. (E) Hematoxylin and eosin (H&E)-stained male heart sections 9 weeks post-TAC. (F) Cardiomyocyte cross-sectional area and width quantified from H&E images (n=3 males, n=3 females per genotype). Scale bar = 30 µm. (G) Periodic acid Schiff-stained male heart sections 9 weeks post-TAC. Arrowheads indicate glycogen aggregates. Scale bar = 35 µm. (H) Masson’s Trichrome-stained male heart sections 9 weeks post-TAC. Arrowheads indicate interstitial collagen. Scale bar = 45 µm. (I) Cardiac fibrosis quantified from Masson’s Trichrome images (n=3 males, n=3 females per genotype). (J) Ejection fraction assessed by echocardiography. (K) Lung weight to body weight ratio assessing pulmonary edema 9 weeks post-TAC (WT: n=3 males, n=5 females; HOIL-1^null^: n=3 males, n=7 females).

### HOIL-1 promotes glycogenolysis

We sought to explore the role of HOIL-1 in cardiomyocyte function in more mechanistic detail. To this end, we modeled the two main PGBM1 genotypes, namely complete knockout or deletion of the C-terminal RBR, using exon-specific gRNA in AC16 human immortalized cardiomyocytes (Figure S1A). Targeting exon 2 of *RBCK1* effectively reduced HOIL-1 protein levels, resulting in destabilized HOIP, the catalytic center of LUBAC, as expected^31^. Exon 7-directed gRNA also reduced full-length HOIL-1, but HOIP levels remained unchanged (Figure S1B). This is consistent with the expression of the N-terminal LUBAC tethering motif (LTM) and ubiquitin-like (UBL) domains in HOIL-1 which stabilize the complex^31^ (Figure S1B). Truncated HOIL-1 had greatly reduced expression by immunoblot, as observed with ΔRING1 HOIL-1 in mouse embryonic fibroblasts (MEFs)^32^. Next-generation sequencing confirmed a frameshift after exon 7, resulting in a premature stop codon (Figure S1C). We hereafter refer to these genotypes as HOIL-1^KO^ (exon 2) and HOIL-1^ΔRBR^ (exon 7).

After confirming HOIL-1^null^ mice accumulate myocardial glycogen aggregates (Figure 1G), we measured glycogen levels in our cell lines. In the heart, glycogen is not used as a primary fuel source, but is rapidly metabolized with increased energetic demand or metabolic stress^33^. We found that during acute glucose starvation, a trigger for glycogenolysis, HOIL-1^ΔRBR^ cells retained more glycogen than both parental and HOIL-1^KO^ cells (Figure S1D). Using LC-MS/MS, we measured a pronounced increase in glucose-1-phosphate, the product of glycogenolysis, in parental and HOIL-1^KO^ cells following acute starvation (Figure S1E). No glucose-1-phosphate was measured in starved HOIL-1^ΔRBR^ cells (Figure S1E), demonstrating that the HOIL-1 RBR domain is necessary for efficient glycogen breakdown during acute nutrient stress.

### The HOIL-1 RBR mitigates nutrient stress

Severe nutrient stress triggers programmed cell death^34^. Given that cardiomyocyte cell death is a hallmark of dilated cardiomyopathy and heart failure^35–38^ and that HOIL-1 has an established role in cell death^39^, we wanted to explore how HOIL-1 loss would impact nutrient stress-induced cell death.

We found that HOIL-1^ΔRBR^ AC16 cells displayed a striking sensitivity to glucose starvation-induced cell death, while complete HOIL-1 loss had no effect (Figure 2A). 786-O renal cell carcinoma are glycogen-rich cells that are resistant to glucose starvation^40^; we generated HOIL-1^ΔRBR^ 786-O renal carcinoma cells and found that they also were hypersensitive (Figure 2B and S1F). HOIL-1^KO^ 786-O cells failed to grow, consistent with a previous report^41^. HOIL-1^ΔRING1^ MEFs, which lack RING1 of the RBR domain^32^ likewise succumbed to glucose starvation more readily than wildtype (Figure 2C and S1G).

**Figure 2.**
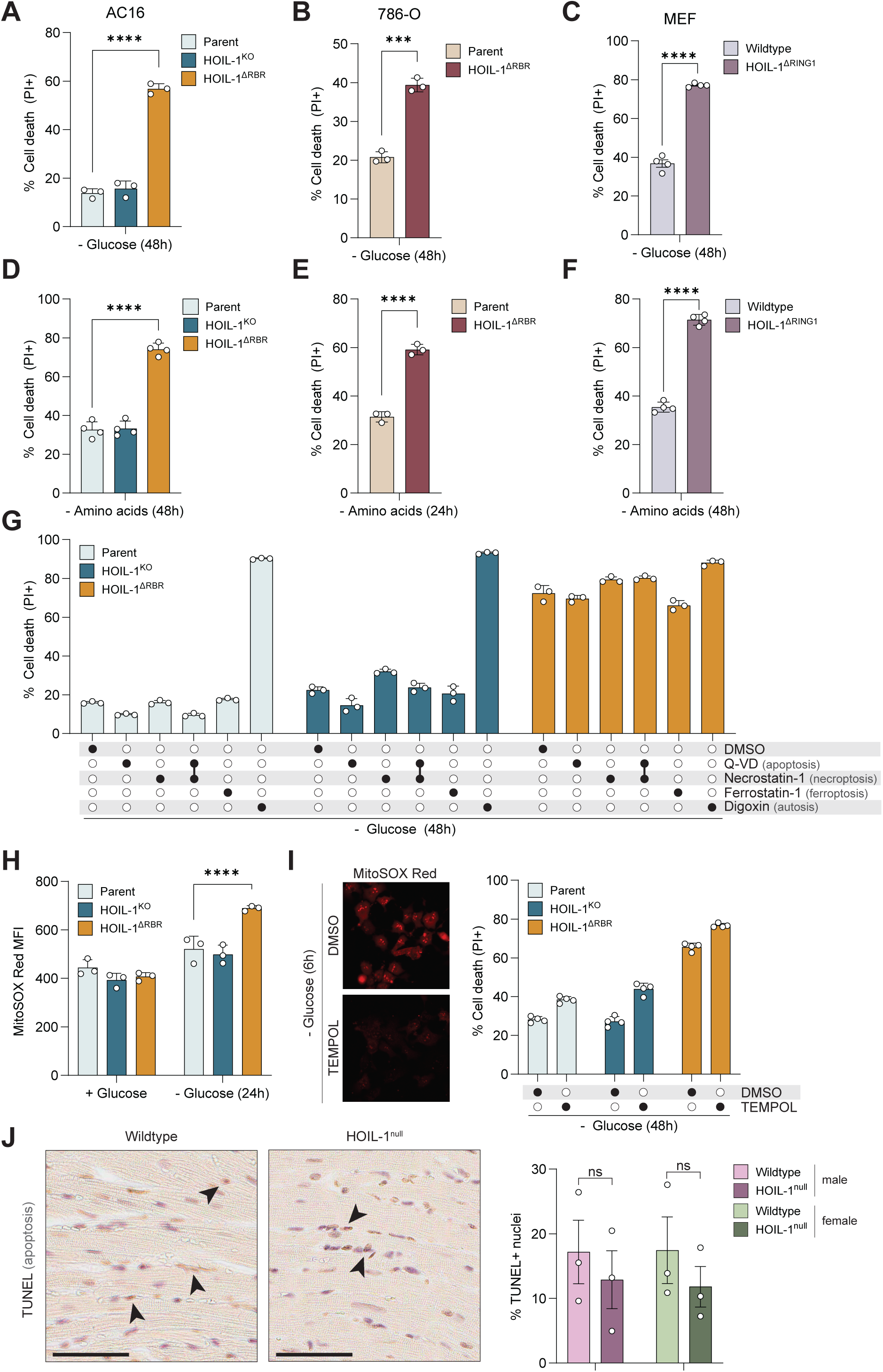
HOIL-1 mitigates nutrient stress. (A) Cell death in AC16 cells after 48h glucose starvation (n=3). (B) Cell death in 786-O cells after 48h glucose starvation (n=3). (C) Cell death in MEFs after 48h glucose starvation (n=4). (D) Cell death in AC16 cells after 48h amino acid starvation (n=4). (E) Cell death in 786-O cells after 24h amino acid starvation (n=3). (F) Cell death in MEFs after 48h amino acid starvation (n=4). (G) Cell death in AC16 cells treated with cell death inhibitors (Q-VD = 5 µM, Necrostatin-1 = 30 µM, Ferrostatin-1 = 2 µM, digoxin = 1 µg/mL) concurrent with 48h glucose starvation (n=3). (H) MitoSOX Red mean fluorescence intensity (MFI) assessed by flow cytometry in AC16 cells grown in full media or after 24h glucose starvation (n=3) (I) Cell death in AC16 cells treated with TEMPOL (50µM) concurrent with 48h glucose starvation (n=3). Representative image of MitoSOX Red staining after TEMPOL shown. (J) TUNEL-stained heart sections 9 weeks post-TAC. Arrowheads indicate TUNEL+ nuclei. Scale bar = 60 µm. Proportion of TUNEL+ nuclei quantified from TUNEL images (n=3 males, n=3 females per genotype, ns= not significant). See also Figures S1, S2

This sensitivity may be attributed to an inability to utilize glycogen. Inhibition of glycogen phosphorylase increased glycogen retention but did not influence cell death in parental or HOIL-1^KO^ AC16 cells (Figure S2A – S2C). Depleting glycogen pools via knockdown of glycogen synthase prior to starvation sensitized AC16 cells to glucose starvation (Figure S2D). Having no accessible glycogen may therefore sensitize cells to glucose starvation-induced cell death.

We next examined the role of other LUBAC components in nutrient stress. CRISPR/Cas9-mediated knockout of HOIP did not affect glucose starvation-induced cell death (Figure S2E). HOIL-1 activity limits LUBAC via autoubiquitination^32^; therefore, LUBAC hyperactivity in HOIL-1^ΔRBR^ cells may drive cell death. Pharmacological inhibition of HOIP using HOIPIN-8 (ref 42) did not protect HOIL-1^ΔRBR^ cells from glucose starvation-induced cell death (Figure S2F). Conversely, and consistent with enhanced LUBAC activity^32^, HOIL-1^ΔRBR^ cells were protected against TNFα-induced apoptosis which could be reversed with HOIPIN-8 (Figure S2F). Finally, while LUBAC hyperactivation also occurs following loss of the linear-specific deubiquitinase OTULIN in some cell types^43,44^, OTULIN silencing had no effect on glucose starvation-induced cell death (Figure S2G). HOIL-1 therefore acts independently of LUBAC in this context.

To identify whether HOIL-1 is more broadly involved in nutrient stress, we starved cells of amino acids and found that HOIL-1^ΔRBR^ AC16 cells, HOIL-1^ΔRBR^ 786-O cells, and HOIL-1^ΔRING1^ MEFs were more susceptible to cell death (Figure 2D-2F). These data demonstrate that the HOIL-1 RBR domain mitigates nutrient stress in multiple cell types.

### Glucose starvation induces noncanonical cell death

Unlike the largely universal apoptotic signaling cascade initiated by death receptor ligands, the mechanism of glucose starvation-induced cell death is context-specific and has been attributed to apoptosis^45–47^, necrosis^48^, necroptosis^49,50^, entosis^51^, and more recently “disulfidptosis”^40,52^.

We sought to characterize the modality of glucose starvation-induced cell death. We tested a panel of inhibitors targeting apoptosis (Q-VD), necroptosis (necrostatin-1), ferroptosis (ferrostatin-1), or autosis (digoxin) and found none prevented cell death in AC16 cells (Figure 2G). Though we found HOIL-1^ΔRBR^ cells had elevated mitochondrial superoxide during glucose starvation, (Figure 2H), the reactive oxygen scavenger TEMPOL had no effect (Figure 2I). We did not observe differential activation of the AMPK and mTOR nutrient sensing pathways by immunoblot, nor could we rescue glucose starvation-induced cell death with pharmacological inhibition or activation of either pathway (Figure S2H and S2I). These data suggest HOIL-1^ΔRBR^ AC16 cells die via a noncanonical cell death program.

Cardiomyocyte cell death precedes heart failure. We next used terminal deoxynucleotidyl transferase dUTP nick end labeling (TUNEL) to assess apoptosis in heart sections from mice that underwent TAC surgery. We found that the extent of apoptosis was variable among mice nine weeks post-surgery and there was no difference between genotypes regardless of sex (Figure 2J). These data are consistent with the observation that HOIL-1^ΔRBR^ cells are protected against classical apoptosis (Figure S2F). Apoptotic cell death therefore does not contribute to the exacerbated cardiomyopathy of HOIL-1^null^ mice.

### HOIL-1 loss alters the nutrient stress metabolome

We reasoned that the noncanonical cell death trigged by glucose starvation may be driven by specific metabolites. We next examined how HOIL-1 deficiency influences the metabolome during nutrient stress. We performed global metabolomics using LC-MS/MS on parental, HOIL-1^KO^ and HOIL-1^ΔRBR^ AC16 cells in glucose-replete media or following acute (3h) or prolonged (24h) glucose starvation (Figure 3A and Table S1). Partial least-squares discriminant analysis (PLS-DA) revealed glucose starvation markedly altered the cellular metabolic profile (Figure 3B). During glucose starvation, HOIL-1^ΔRBR^ cells at both timepoints clustered more closely with parent AC16 cells than HOIL-1^KO^ cells (Figure 3B). These data demonstrate that HOIL-1 loss alters the nutrient stress metabolic signature.

**Figure 3.**
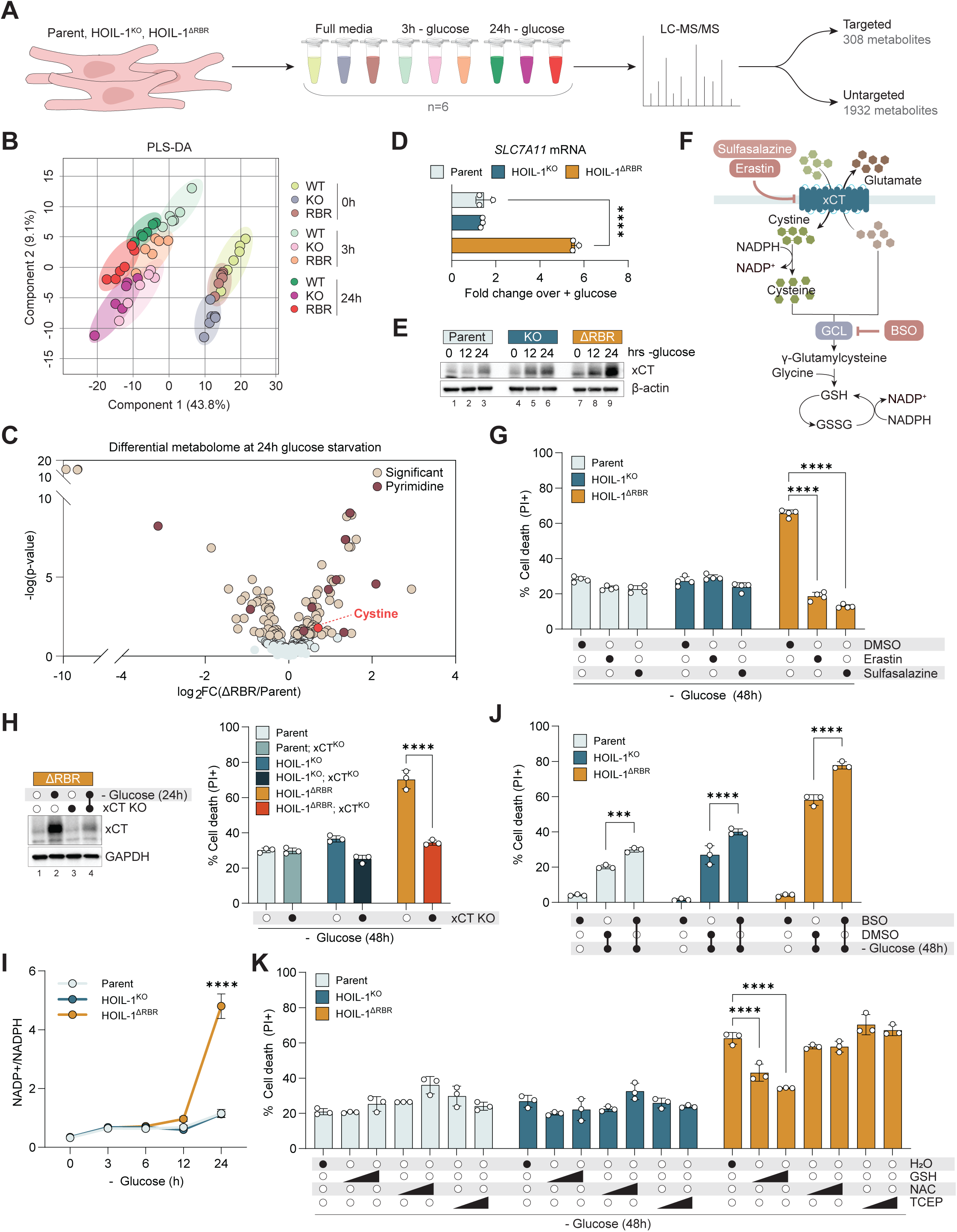
The nutrient stress metabolome reveals glucose starvation causes noncanonical cell death mediated by cystine influx via xCT. (A) Schematic of the experimental design examining the metabolome in AC16 cells in full media or after 3h or 24h glucose starvation. (B) Partial least squares discriminant analysis (PLS-DA) of the metabolomics dataset. (C) Volcano plot of targeted metabolites in parent and HOIL-1^ΔRBR^ AC16 cells after 24h glucose starvation. Significant: −log(p-value) ≥1.3. (D) *SLC7A11* mRNA levels (normalized to *TBP*) in AC16 cells after 24h glucose starvation. Data represented as the fold change over full media (n=3). (E) Immunoblot analysis of xCT (SLC7A11) levels in AC16 cells during glucose starvation. (F) Schematic of xCT-dependent nutrient and metabolite flux. (G) Cell death in AC16 cells treated with erastin (2 µM), sulfasalazine (500µM), or DMSO vehicle concurrent with 48h glucose starvation (n=4). (H) Cell death in xCT^KO^ AC16 cells after 48h glucose starvation (n=3). Representative immunoblot of xCT knockout shown. (I) Measurement of NADP+ and NADPH ratios during glucose starvation in AC16 cells (n=3). (J) Cell death in AC16 cells treated with BSO (500 µM) or DMSO vehicle concurrent with 48h glucose starvation (n=3). (K) Cell death in AC16 cells treated with GSH ethyl ester (0.5 mM, 1.0 mM), NAC (0.5 mM, 1.0 mM), TCEP (0.5 mM, 1.0 mM), or H_2_O vehicle concurrent with 48h glucose starvation (n=3). See also: Figures S3, S4

### xCT promotes cell death by disrupting glutathione homeostasis

We focused on metabolites with increased abundance in HOIL-1^ΔRBR^ cells during prolonged starvation as putative drivers of cell death. Pyrimidines were among the most enriched metabolites in starved HOIL-1^ΔRBR^ cells relative to parent cells (Figure 3C, S3A, and S3B), so we assessed their contribution. Hydroxyurea (a ribonucleotide reductase inhibitor), brequinar (an inhibitor of *de novo* pyrimidine synthesis), and oxythiamine (a transketolase inhibitor) all failed to rescue cell death (Figure S3C and S3D). Likewise, supplemental uridine did not sensitize cells to glucose starvation (Figure S3D).

Cystine was among the most differentially abundant metabolites in HOIL-1^ΔRBR^ cells (Figure 3C). Cystine is imported into the cytosol via system xc^-^, a cystine-glutamate antiporter consisting of SLC7A11, also known as xCT, and SLC3A2 that previously has been implicated in glucose starvation-induced cell death^40,52^. Consistent with increased cystine, we observed robust upregulation at the mRNA (Figure 3D) and protein (Figure 3E) levels in HOIL-1^ΔRBR^ cells during prolonged glucose starvation. xCT inhibition using either erastin or sulfasalazine (Figure 3F and 3G) and CRISPR/Cas9-mediated xCT knockout (Figure 3H) both substantially blocked cell death. Conversely, stable xCT overexpression rendered parental AC16 cells sensitive to glucose starvation, which could be blocked by erastin (Figure S3E). Glucose starvation-induced cell death was partially rescued by depleting media of cystine (Figure S3F), demonstrating that xCT drives the distinct cell death observed in HOIL-1^ΔRBR^ cells in part via cystine import.

Cystine is highly insoluble^53^ and is rapidly reduced to two molecules of cysteine in the cytosol using NADPH. The pentose phosphate pathway (PPP) generates the bulk of cellular NADPH from glucose-6-phosphate (Figure S4A). xCT activity during glucose starvation therefore consumes NADPH without replenishing it, increasing the ratio of NADP+/NADPH. We indeed found a sharp increase in the NADP+/NADPH ratio in glucose-starved HOIL-1^ΔRBR^ cells (Figure 3I), which was normalized to parental levels by knocking out xCT (Figure S4B). We interrogated whether collapse of the NADPH system was involved in cell death, as reported in other systems^54^. Inhibiting glucose-6-phosphate dehydrogenase (G6PD), the rate-limiting enzyme of the PPP, using G6PDi-1^55^ increased the NADP+/NADPH ratio (Figure S4C) but did not sensitize cells to glucose starvation (Figure S4D). 6-aminonicotinamide, another G6PD inhibitor, also had no effect (Figure S4D). Next, we generated cells with doxycycline-inducible expression of the engineered bacterial oxidase TPNOX, which oxidizes NADPH orthogonal to mammalian cell biology^56^. No effect on cell death was observed whether TPNOX was induced prior to starvation or concurrently (Figure S4E). These data suggest that NADPH collapse *per se* is not causing cell death.

Increased NADP+/NADPH ratios reflect a deficit in cellular reducing power. This was supported by the accumulation of the glutathione (GSH) precursor γ-glutamylcysteine and oxidized GSH (GSSG) in HOIL-1^ΔRBR^ cells as detected in our untargeted metabolomics screen (Figure S4F). We found that GSH synthesis inhibition with buthionine sulfoximine (BSO) enhanced glucose starvation-induced cell death (Figure 3J). We next assessed the effect of exogenous reducing agents. While neither N-acetylcysteine nor tris(2-carboxyethyl)phosphine (TCEP) had a protective effect, GSH ethyl ester dose-dependently suppressed glucose starvation-induced cell death (Figure 3K). These data collectively show that selective upregulation of xCT in HOIL-1^ΔRBR^ cells during glucose starvation drives cell death by disrupting GSH homeostasis.

### The integrated stress response effector ATF4 orchestrates cell death

Why is xCT uniquely induced in HOIL-1^ΔRBR^ cells during glucose starvation? xCT is a target gene of ATF4 and NRF2, two key transcription factors in the ISR and oxidative stress response, respectively. We found that both ATF4 and NRF2 were selectively upregulated in HOIL-1^ΔRBR^ AC16 cells during starvation (Figure 4A). There was a pronounced induction of ATF4 and, to a lesser extent, NRF2 target genes in HOIL-1^ΔRBR^ cells (Figure 4B), further demonstrating activation of these transcription factors. Knockdown of ATF4, but not NRF2, abolished xCT upregulation and subsequent cell death (Figure 4C and 4D). We observed that ATF4 knockdown also suppressed NRF2: this regulation was post-transcriptional, specific to glucose starvation, and independent of the GSH-degrading enzyme CHAC1 (Figure S5A - S5C), which was recently identified as a link between these two proteins^57^. Curiously, eIF2α phosphorylation, which represses global translation while driving ATF4 translation, was only slightly increased in HOIL-1^ΔRBR^ cells (Figure 4A). This accompanied a modest decrease in global protein synthesis in HOIL-1^ΔRBR^ cells relative to parental and HOIL-1^KO^ cells as assessed by puromycin incorporation (Figure S4D). We further found no difference between cell lines in either BiP induction, a marker of endoplasmic reticulum (ER) stress, or cleavage of PGAM5, a mitochondrial phosphatase that is cleaved following loss of mitochondrial membrane potential^58^ (Figure 4A). These data reflect an equivalent initial stress burden from glucose starvation across our cell lines. ATF4 translation therefore appears divorced from the canonical upstream pathway in this context. Glucose starvation-induced cell death was also driven by ATF4 and xCT in HOIL-1^ΔRBR^ 786-O cells (Figure 4E – 4G).

**Figure 4.**
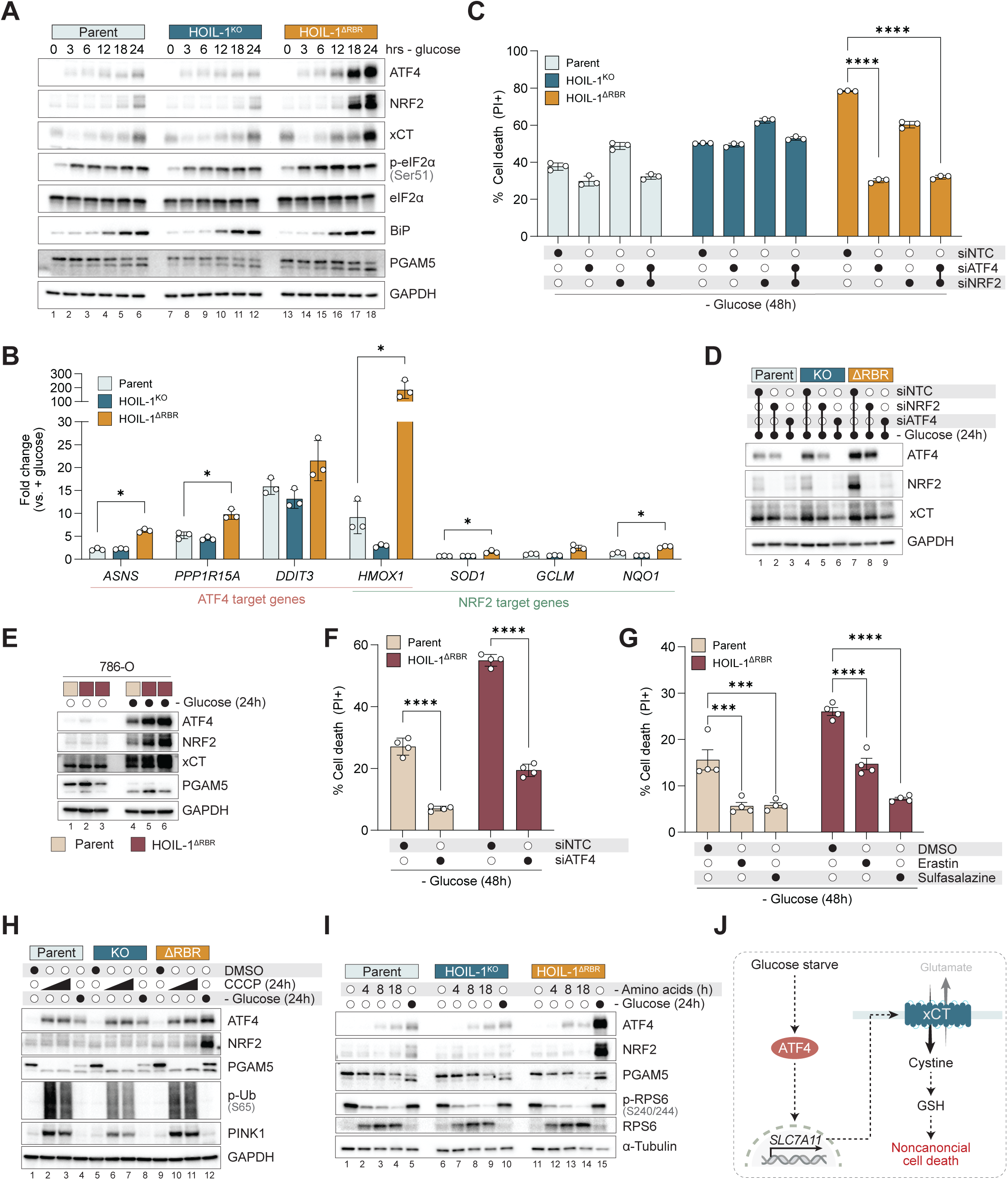
ATF4 orchestrates cell death. (A) Immunoblot analysis of AC16 cells after glucose starvation for the indicated time-points. (B) ATF4 and NRF2 target genes normalized to *TBP* in AC16 cells measured by RT-qPCR. Data represented as the fold change over full media (n=3). (C) Cell death in AC16 cells transfected with ATF4, NRF2, or non-targeting control (NTC) siRNA 48h prior to glucose starvation for 48h (n=3). (D) Immunoblot analysis of AC16 cells transfected with ATF4, NRF2, or NTC siRNA 48h prior to glucose starvation for 24h. (E) Immunoblot analysis of 786-O cells after 24h glucose starvation. Two different HOIL-1^ΔRBR^ clones shown. (F) Cell death in 786-O cells transfected with ATF4 or NTC siRNA 48h prior to glucose starvation for 48h (n=4). (G) Cell death in 786-O cells treated with erastin (2 µM), sulfasalazine (500 µM), or DMSO vehicle concurrent with 48h glucose starvation (n=4). (H) Immunoblot analysis of AC16 cells after DMSO vehicle, CCCP (5 µM, 25 µM), or glucose starvation for 24h. (I) Immunoblot analysis of AC16 cells after amino acid or glucose starvation for the indicated time-points. (J) Schematic of cell death pathway. See also: Figure S5

We next examined whether HOIL-1^ΔRBR^ cells preferentially engage ATF4 in response to other metabolic stressors and found that neither mitochondrial oxidative phosphorylation uncoupling with cyanide m-chlorophenylhydrazone (CCCP) nor amino acid starvation induced a robust ISR (Figure 4H and 4I).

These data collectively demonstrate that loss of the HOIL-1 RBR domain results in maladaptive activation of the ISR effector ATF4 during glucose starvation, leading to xCT-dependent cell death (Figure 4J).

### Selective autophagy is not involved in the glucose starvation response

Since the RBR of HOIL-1 confers E3 ligase activity, we explored global changes to the ubiquitinome during glucose starvation. We performed ubiquitin remnant profiling prior to the onset of the ISR to identify molecular events upstream of ATF4 (Figure 5A and S6A). After 8 hours of starvation, there were 186 upregulated and 321 downregulated ubiquitinated peptides (log2FC > 1.04 or < −1.04, FDR < 0.05) in HOIL-1^ΔRBR^ cells relative to parental AC16 cells (Table S2).

**Figure 5.**
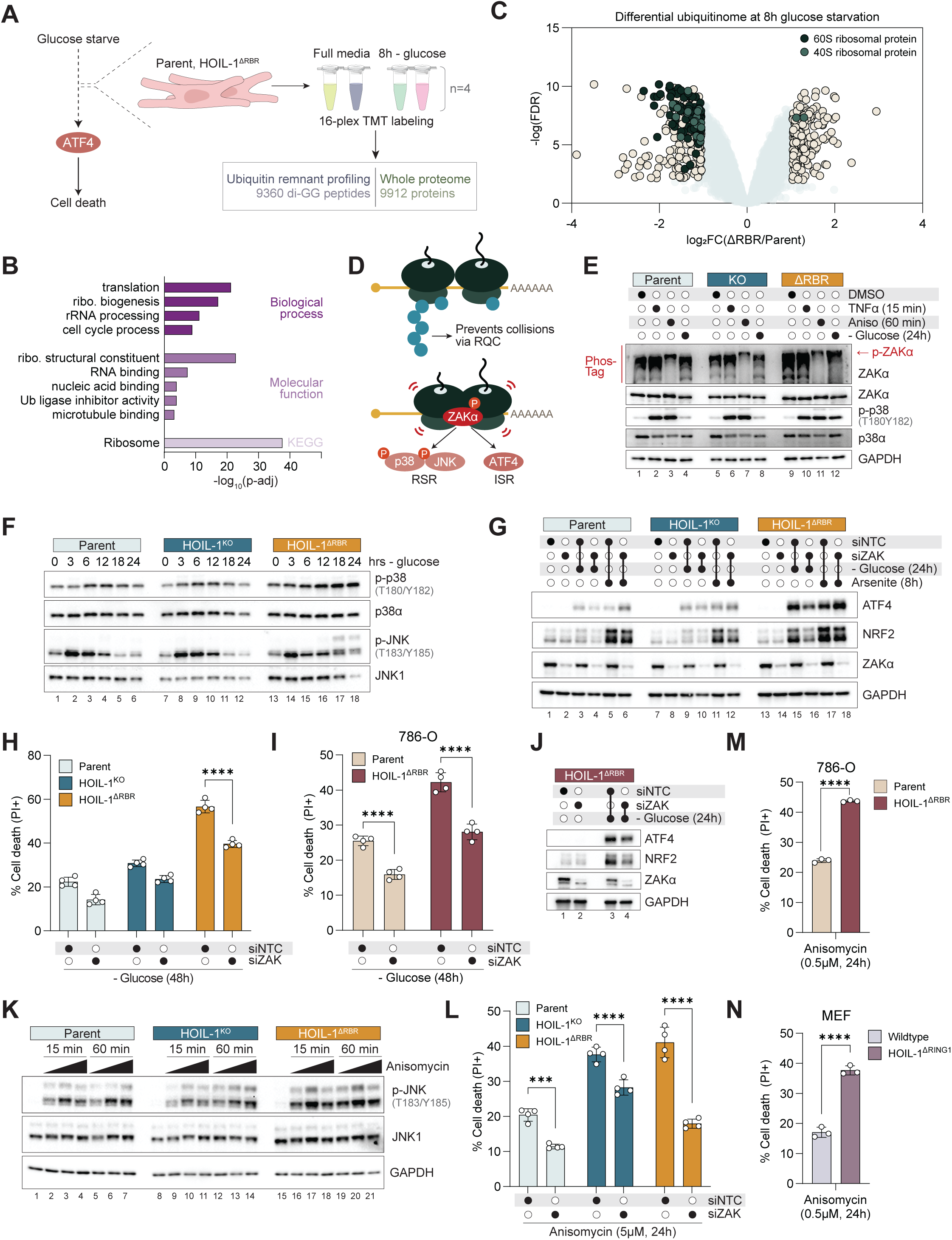
Loss of the HOIL-1 RBR domain remodels the ribosome ubiquitin landscape and sensitizes cells to ZAKα-activating ribotoxic stressors. (A) Schematic of the experimental design examining the ubiquitinome in AC16 cells in full media or after 8h glucose starvation. (B) Gene ontology enrichment analysis of proteins with reduced ubiquitin-modified peptides in HOIL-1^ΔRBR^ cells relative to parent AC16 cells after 8h glucose starvation. (C) Volcano plot showing differentially ubiquitinated peptides after 8h glucose starvation. Significant: −log(FDR) ≥ 2SD, ≤ −2SD. (D) Schematic of ribosomal stress signaling. Ubiquitination of stalled ribosomes promotes their extraction via ribosome quality control (RQC) to prevent collisions. Ribosome collisions activate ZAKα which engages the ribotoxic stress response (RSR) via the MAPKs p38 and JNK and integrated stress response (ISR) via ATF4. (E) Immunoblot analysis of AC16 cells after DMSO vehicle, TNFα (10 ng/mL), anisomycin (5 µM), or glucose starvation for the indicated time-points. ZAKα phosphorylation was assessed using a Phos-Tag acrylamide gel. (F) Immunoblot analysis of AC16 cells after glucose starvation for the indicated time-points. (G) Immunoblot analysis of AC16 cells transfected with ZAK (targeting ZAKα and ZAKβ) or non-targeting control (NTC) siRNA 48h prior to sodium arsenite (10 µM) or glucose starvation for the indicated time-points. (H) Cell death in AC16 cells transfected with ZAK or NTC siRNA 48h prior to glucose starvation for 48h (n=4). (I) Cell death in 786-O cells transfected with ZAK or NTC siRNA 48h prior to glucose starvation for 48h (n=4). (J) Immunoblot analysis of 786-O cells transfected with ZAK or NTC siRNA 48h prior to glucose starvation for 24h. (K) Cell death in AC16 cells transfected with ZAK or NTC siRNA 48h prior to anisomycin (5µM) treatment for 24h (n=4). (L) Immunoblot analysis of AC16 cells after low (0.1 µM), medium (5 µM), and high (250 µM) dose anisomycin treatment for the indicated time-points. (M) Cell death in 786-O cells after 24h anisomycin treatment (0.5 µM) (n=3). (N) Cell death in MEFs after 24h anisomycin treatment (0.5 µM) (n=3). See also: Figures S6, S7

Gene ontology analysis of proteins with increased ubiquitination revealed an overrepresentation of chaperones and proteins regulating stress responses and protein processing in the ER (Figure S6B). Ubiquitination and chaperones play a critical role in selective autophagy of the ER (ER-phagy); indeed, there were data suggesting increased ER-phagy activity. HOIL-1^ΔRBR^ cells had increased ubiquitinated ARL6IP1 (Figure S6C), an ER-shaping protein whose ubiquitination was recently shown to promote ER-phagy flux^59^. Further, whole proteome analysis revealed that the level of collagens, a prototypical ER-phagy substrate^60^, was dramatically reduced in HOIL-1^ΔRBR^ cells (Table S3 and Figure S6D), which we confirmed by western blot (Figure S6E). Nevertheless, silencing ARL6IP1 or the ER-phagy receptor FAM134B did not reduce ATF4 translation (Figure S6F). Unlike amino acid starvation or mTOR inhibition, glucose starvation failed to induce ER-phagy flux using two different reporters^61,62^ (Figure S6G - S6I).

Gene ontology analysis of proteins with decreased ubiquitination revealed a significant enrichment of ribosome-associated proteins and processes (Figure 5B). Indeed, the most striking feature of the ubiquitinome dataset was the dramatic reduction in both 40S and 60S ribosomal protein ubiquitination in HOIL-1^ΔRBR^ cells (Figure 5C), prompting us to explore ribophagy. There was no change in the abundance of ribosomal proteins (Figure S6E), nor did we detect ribophagy flux using a pH-sensitive reporter^63^ (Figure S6J and S6K). Collectively, these data suggest selective autophagy is not involved in the glucose starvation response.

### The ribosome collision sensor ZAKα activates ATF4 during nutrient stress

Ubiquitination also plays a central role in ribosome quality control (RQC) (Figure 5D). Under conditions of translational stress, a ribosome may stall on its cognate mRNA, leading to a collision by a trailing ribosome. Ubiquitination of stalled ribosomes facilitates their extraction from the mRNA to prevent further collisions^64,65^. As such, inadequate RQC leads to excessive ribosome collisions^66,67^. These collisions are sensed by the MAP3K ZAKα, which autophosphorylates at the interface of two collided ribosomes to engage a two-pronged stress response involving phosphorylation of stress-activated protein kinases p38 and JNK via the ribotoxic stress response (RSR) and the ISR via GCN1/GCN2-mediated eIF2α phosphorylation^2^.

We examined ZAKα activation as a marker of ribosomal stress. During glucose starvation, ZAKα and its downstream target p38 were phosphorylated uniquely in HOIL-1^ΔRBR^ cells (Figure 5E). Conversely, the translation elongation inhibitor anisomycin, a known inducer of ribosome collisions, triggered ZAKα activation in all cell lines. A time-course analysis demonstrated that p38 phosphorylation occurred prior to ATF4 induction (Figure 5F). ZAK silencing, which targets both ZAKα and ZAKβ isoforms, attenuated ATF4 in response to glucose starvation but not sodium arsenite, a general inducer of oxidative stress (Figure 5G) and protected against cell death in HOIL-1^ΔRBR^ AC16 and 786-O cells (Figure 5H – 5J). ZAKα-driven ATF4 upregulation during glucose starvation was independent of GCN1 and GCN2; in fact, GCN2 silencing markedly enhanced ATF4 and cell death (Figure S7A and S7B).

We next investigated whether HOIL-1 RBR domain loss sensitized cells to other ribotoxic stressors. Following anisomycin treatment, JNK phosphorylation and cell death were enhanced in HOIL-1^ΔRBR^ AC16 cells, which was mediated by ZAKα (Figure 5K and 5L). HOIL-1^ΔRBR^ 786-O and HOIL-1^ΔRING1^ MEFs were also hypersensitive to anisomycin-induced cell death (Figure 5M and 5N), demonstrating a broader role for HOIL-1 in restricting the RSR.

### Glucose starvation activates ZAKα in HOIL-1^ΔRBR^ cells through a unique mechanism

ZAKα is heretofore known to be activated only by ribosome collisions. After obtaining several lines of evidence that ZAKα is activated in HOIL-1^ΔRBR^ cells, we performed sucrose gradient fractionation to examine the re-localization of collision-associated proteins to heavier ribosome fractions following anisomycin treatment or glucose starvation (Figure S7C - S7F). Higher order ribosome complexes accumulated more in HOIL-1^ΔRBR^ cells than parental cells following anisomycin treatment (Figure S7D), which is consistent with more collisions^68^. Further, in response to anisomycin, we saw a clear shift in EDF1, a collision-specific marker^69^, by immunoblot analysis of sucrose fractions (Figure S7F). In contrast, glucose starvation triggered the collapse of polysomes into 80S monosomes (Figure S7E). Glucose starvation had no effect on EDF1 or GIGYF2, another collision marker^70,71^ (Figure S7F). We also observed no differential shift in ZAKα between genotypes. These data may reflect a distinct collision assembly during glucose starvation, a collision-independent mode of ZAKα activation, or a technical challenge.

Ribosome collisions may be the result of excessive stalling rather than defective quality control. We examined the effect of HOIL-1 loss on ribosome stalling using a dual fluorescence reporter consisting of GFP and RFP separated by a linker of 20 lysines encoded by AAA codons, or a Flag tag as a non-stalling control^72^ (Figure S7G). Poly(A) stretches cue ribosome stalling^73^, impairing downstream RFP translation. As a control, silencing of ZNF598, a RQC E3 ligase that promotes stalling^64^, increased the RFP:GFP ratio (Figure S7H). HOIL-1^KO^ cells had a modest defect in stalling, while HOIL-1^ΔRBR^ cells did not (Figure S7D).

Collectively, these data identify the HOIL-1 RBR domain as an engineer of the ribosome ubiquitin landscape that controls cell fate under translational stress independent of ribosome stalling. Failure of this system engages a ZAKα-dependent cell death.

### HOIL-1 proximitome during glucose starvation

How could loss of the HOIL-1 RBR cause widespread changes to ribosome ubiquitination? To address this question, we sought to characterize the HOIL-1 proximal network during glucose starvation using proximity labeling. We fused the MiniTurbo biotin ligase at either the N- or C-terminus of ΔRBR HOIL-1 and stably expressed these constructs in HOIL-1^ΔRBR^ AC16 cells, using GFP-MiniTurbo as a negative control (Figure 6A). This approach identified 24 proteins across two experiments that were enriched in both HOIL-1-MiniTurbo cell lines relative to GFP-MiniTurbo after 6 hours of starvation, of which, unexpectedly, half (12/24) were RNA-binding proteins (Table S4 and Figure 6B). Among these 24, only PABPC1 had decreased ubiquitination in HOIL-1^ΔRBR^ cells during glucose starvation (Figure S8A), highlighting its potential as a direct HOIL-1 substrate. STRING analysis identified PABPC1 as a hub of the HOIL-1 proximitome, further supporting its candidacy (Figure S8B).

**Figure 6.**
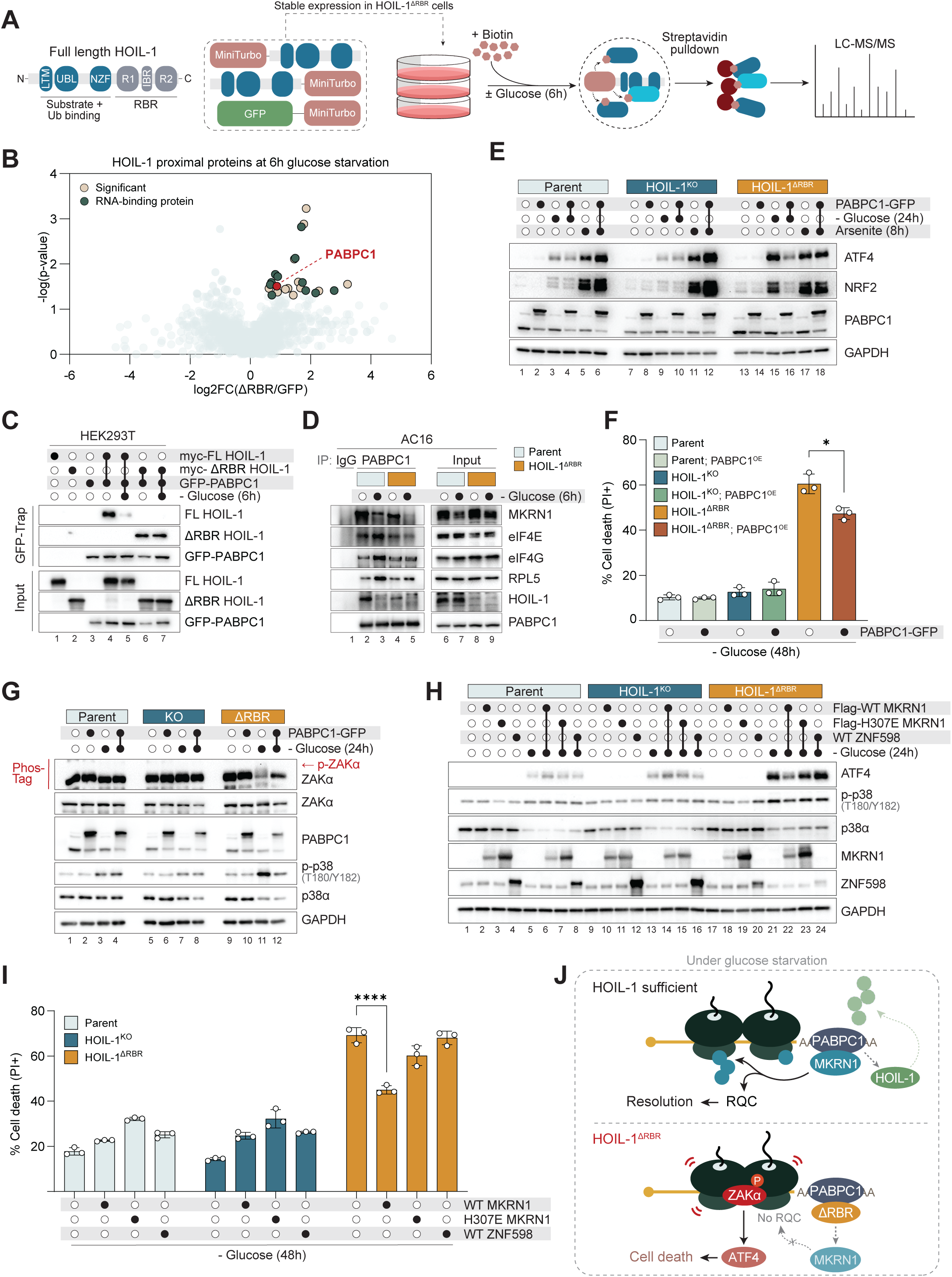
Mutant HOIL-1 locks onto PABPC1 and excludes MKRN1, rendering glucose starvation ribotoxic. (A) Schematic of the experimental design examining the ΔRBR HOIL-1 proximitome in AC16 cells in full media or after 6h glucose starvation. (B) Volcano plot showing log2 fold changes of proteins detected in ΔRBR HOIL-1-MiniTurbo streptavidin pulldown samples over GFP-MiniTurbo samples after 6h glucose starvation. Significant: −log_10_(p-value)>1.3, log_2_FC>0.6 (C) Immunoblot analysis of HEK293T cells transfected with the indicated plasmids 24h prior to 6h glucose starvation where indicated, followed by lysis and GFP-Trap immunoprecipitation. (D) Immunoblot analysis following endogenous PABPC1 or IgG immunoprecipitation from AC16 lysates. (E) Immunoblot analysis of AC16 cells stably expressing PABPC1-GFP following sodium arsenite treatment (10 µM) for 8h or glucose starvation for 24h. (F) Cell death in AC16 cells stably expressing PABPC1-GFP after 48h glucose starvation (n=3). (G) Immunoblot analysis of AC16 cells stably expressing PABPC1-GFP after 24h glucose starvation. ZAKα phosphorylation was assessed using a Phos-Tag acrylamide gel. (H) Immunoblot analysis of AC16 cells stably expressing MKRN1 or ZNF598 after 24h glucose. (I) Cell death in AC16 cells stably expressing MKRN1 or ZNF598 after 48h glucose starvation (n=3). (J) Model of HOIL-1 activity in ribosome stress signaling. See also: Figure S8

### Mutant HOIL-1 locks onto PABPC1 and excludes MKRN1, rendering glucose starvation ribotoxic

PABPC1 is a cytosolic poly(A) binding protein whose ability to regulate virtually all aspects of mRNA metabolism is dictated by the various protein complexes it assembles. For example, PABPC1 promotes translation initiation via mRNA circularization through interaction with the eIF4E/eIF4G cap-binding complex^74,75^, stimulates deadenylation and prevents mRNA decay via interaction with the CCR4-NOT complex^76,77^, and, critically, promotes ribosome ubiquitination via interaction with Makorin (MKRN) E3 ligases^78^.

We first validated the interaction between PABPC1 and HOIL-1 in HEK293T cells. Full-length HOIL-1 bound PABPC1 in glucose-replete media but dissociated during glucose starvation (Figure 6C). The interaction between ΔRBR HOIL-1 and PABPC1 was weaker than full-length but did not change during glucose starvation (Figure 6C). We found that co-expression of full-length HOIL-1 triggered PABPC1 ubiquitination, which decreased during starvation (Figure S8C). As expected, ΔRBR HOIL-1 had no effect on ubiquitination, while MKRN1, a known PABPC1 E3 ligase^78^, robustly modified PABPC1 (Figure S8C).

One outstanding question from our data thus far is: what explains the disparity in phenotypes between HOIL-1^KO^ and HOIL-1^ΔRBR^ cells? One possibility is that truncated HOIL-1 acts as a dominant negative, wherein it binds to its substrate but does not ubiquitinate or dissociate from it. We therefore examined PABPC1 interactors during glucose starvation in HOIL-1^ΔRBR^ AC16 cells. As observed in HEK293T cells, full-length HOIL-1 binds endogenous PABPC1 in AC16 cells but dissociates during glucose starvation (Figure 6D). This was concomitant with the recruitment of known PABPC1 binding partners eIF4G, eIF4E, and RPL5 (Figure 6D). This recruitment is abrogated in HOIL-1^ΔRBR^ cells, suggesting truncated HOIL-1 prevents PABPC1 from binding its functional partners.

MKRN1 is recruited to ribosomes via PABPC1, where it initiates RQC by ubiquitinating RPS10, triggering a cascade of downstream ubiquitination events^78^. We found that the interaction between PABPC1 and MKRN1 is abolished in glucose-starved HOIL-1^ΔRBR^ cells (Figure 6D). Subcellular fractionation of RNA-protein granules^79^ further revealed MKRN1 is absent from this fraction in HOIL-1^ΔRBR^ cells during starvation despite equivalent PABPC1 levels (Figure S8D, lanes 46-47). These data suggest that MKRN1 is mislocalized in HOIL-1^ΔRBR^ cells, preventing it from initiating a cytoprotective RQC response.

If HOIL-1^ΔRBR^ is acting as a dominant negative, we reasoned that by altering the stoichiometry of the system we could override its effects. Indeed, PABPC1 overexpression shut down ATF4 in response to glucose starvation but not sodium arsenite (Figure 6E) and protected against glucose starvation-induced cell death (Figure 6F). Ribotoxic stress was quelled by PABPC1 overexpression, as determined by reduced ZAKα and p38 phosphorylation (Figure 6G). Overexpression of wildtype MKRN1, but not the catalytically inactive H307E mutant, likewise suppressed glucose starvation-induced ZAKα activation, ATF4, and cell death (Figure 6H and 6I). Critically, another RQC E3 ligase, ZNF598, could not rescue cell death (Figure 6I), reflecting the importance of MKRN1 E3 ligase activity and the consequences of its mislocalization. Neither PABPC1 nor MKRN1 overexpression rescued amino acid starvation-induced cell death (Figure S8E and S8F), revealing their selective involvement in the glucose starvation response.

These data identify HOIL-1 as an inhibitor of PABPC1 activity. We show that HOIL-1 constitutively binds PABPC1 but dissociates during glucose starvation to facilitate the binding of functional partners, including MKRN1. This allows the engagement of a cytoprotective RQC response and resolution of stress. We find that a disease-relevant mutation of HOIL-1 locks it on PABPC1 and blocks these interactions. This excludes MKRN1 from the complex, thereby limiting RQC and precipitating a ZAKα-ATF4-driven stress response that kills the cell (Figure 6J).

## DISCUSSION

Cardiomyocytes are omnivorous. They are able to adapt their metabolic substrate usage according to energetic demands. Under resting conditions, mitochondrial fatty acid oxidation accounts for most of their cellular ATP production. Cardiac distress, however, is associated with loss of this metabolic flexibility and a shift towards increased reliance on glycogen, glucose, and amino acids^80,81^. Glucose release from selective autophagy of glycogen particles via glycophagy is thought to play an important role in cardiac metabolic stress^82^ and diabetic cardiomyopathy^83^. As HOIL-1 is now known to be able to directly ubiquitinate glycogen *in vitro*^25,84,85^, which we also confirmed (data not shown), a compelling hypothesis that follows is that HOIL-1 promotes glycophagy by tagging glycogen particles for clearance. The inability of cardiomyocytes in HOIL-1^null^ mice to breakdown glycogen, perhaps due to defective glycophagy, may disrupt the adaptive metabolic changes needed to mitigate pressure overload, thereby accelerating pathology (Figure 1). Indeed, impaired glycogen breakdown, as in Danon and Pompe disease^86^, or excessive glycogen deposition, as in PRKAG2 syndrome^87^, manifests as cardiomyopathy. In agreement with recent data^3,4^, we propose that these metabolic perturbations may also cause ribosomal stress.

Male, but not female, HOIL-1^null^ mice had exacerbated cardiomyocyte cellular hypertrophy and LV dilatation following LV pressure overload *in vivo*. While there is no known sex bias in PGBM1^15^, female mice have distinct gene expression profiles in cardiomyocytes following TAC surgery^88^ which may explain their differential response. Notably, ZAK-deficient mice also present sex-biased metabolic phenotypes. Male ZAK^-/-^ mice are slightly leaner than WT mice, while females have similar body mass^3^. Further, male ZAK^-/-^ mice are protected against aging-associated metabolic dysfunction while females are not^4^.

There is prior evidence that ZAKα contributes to cardiac pathophysiology. In fact, ZAKα was first cloned from a cDNA library derived from a failing human heart in search of regulators of cardiac hypertrophy^89^. ZAKα is abundantly expressed in cardiomyocytes^90^ and, when overexpressed, promotes hypertrophy of cardiac myoblasts^91^. Transgenic overexpression of the ZAKβ isoform in cardiomyocytes *in vivo* causes spontaneous cardiac fibrosis and accelerated mortality in response to isoproterenol-induced hypertrophy^92^. Doxorubicin is a chemotherapeutic agent with pronounced cardiotoxic side-effects that was shown to activate the RSR via ZAKα^93,94^. Finally, a selective ZAKα inhibitor protected against hypertrophic cardiomyopathy in a spontaneous hypertensive rat model^95^. Though we lack direct evidence, we speculate that sex-specific ZAKα activity may in part drive cardiac hypertrophy in HOIL-1^null^ mice. Future investigation will address this important question.

Our data support the notion that cells have evolved strategies of mitigating nutrient stress by signaling via the ribosome^2^. We show that glucose starvation by default is not a ribotoxic stressor but can become one in the absence of the HOIL-1 RBR domain (Figure 5E - 5G). We have identified distinct roles for HOIL-1 in promoting glycogen breakdown and facilitating RQC. The defect in both processes in HOIL-1^ΔRBR^ cells during glucose starvation may engender ribosomal stress exceeding the threshold for ZAKα-mediated ISR activation. Ribosomes stall towards the 3’ end of transcripts during glucose starvation in yeast^96^. Conversely, amino acid starvation or oxidative stress leads to stalling towards the start codon^96^. Given that both MKRN1 and PABPC1 reside at the start of poly(A) tails and in A-rich stretches of 3’ UTRs^78^, we speculate that their distinct roles in the glucose starvation response is due to their proximity to the stalled ribosomes.

In our system, glucose starvation causes a noncanonical cell death mediated by the cystine-glutamate antiporter xCT. Despite similar regulators, this modality is distinct from ferroptosis: ferroptosis is caused by xCT inhibition, whereas xCT inhibition potently rescues glucose starvation-induced cell death. Additionally, preventing lipid peroxidation with ferrostatin-1 afforded no protection during glucose starvation, but fully blocked conventional ferroptosis (Figure S3G). Our results corroborate previous work that implicates xCT in noncanonical cell death^40,52,53^. We found that exogenous GSH dose-dependently protected against cell death, while other antioxidants or reducing agents could not. Though we have not identified the precise mechanism by which cells die, we conclude that the accumulation of oxidized GSH, possibly causing excess protein glutathionylation, is specifically implicated. One limitation of our study is that we have not yet established whether this cell death occurs *in vivo*. Understanding the physiological relevance of this noncanonical cell death modality and its contribution to disease will be an important area of investigation. Our findings nonetheless show that loss of the HOIL-1 RBR domain dramatically enhances susceptibility to glucose starvation-induced cell death in multiple cell types.

Ribosomes are massive macromolecular assemblies. Beyond their textbook roles in protein translation, their vast surfaces also serve as signaling platforms where information about the cellular metabolic state can be encoded. The language of ubiquitin is well-suited for this coding. While several of the ribosomal protein ubiquitination sites identified here have described functions, most of them do not yet. Decoding this information and understanding how E3 ligases and deubiquitinases collaborate in this context will be of great interest. Advances in techniques to detect noncanonical ubiquitin modifications, such as HOIL-1-mediated ester-linked ubiquitin, will undoubtedly reveal new layers of regulation of ribosomal stress.

## ACKNOWLEDGEMENTS

This work was supported in part by NIH R35CA197589 to CMC. We thank Rebecca Cardone, Qiushi Sun, and the Chemical Metabolism Core at Yale University for LC-MS/MS global metabolomics; Nicole Guerrera for echocardiography; Jean Kanyo, Florine Collin, and the Keck Proteomics Core at Yale University for LC-MS/MS analysis of proximity labeling experiments. HOIL-1^null^ were kindly provided by Donna MacDuff (University of Illinois at Chicago). The graphical abstract was designed with SciStories LLC.

## AUTHOR CONTRIBUTIONS

Conceptualization, T.D.; Methodology, T.D., J.Z., Z.W., and K.A.; Investigation, T.D., J.Z., Z.W., K.A., and M.M.; Resources, K.I.; Writing – Original Draft, T.D.; Writing – Review & Editing, T.D., L.H.Y., and C.M.C.; Visualization, T.D.; Supervision, T.D., W.V.G., J.P., L.H.Y., and C.M.C.; Project Administration, T.D.; Funding Acquisition, C.M.C..

## DECALARATION OF INTERESTS

The authors declare no competing interests.

## Methods

### RESOURCE AVAILABILITY

#### Lead contact

Further information and requests for reagents and resources should be directed to and will be fulfilled by the lead contact, Todd Douglas.

#### Materials availability

Materials and reagents used in this study are listed in the key resources table. Reagents generated in this study are available upon request.

#### Data and code availability

Raw mass spectrometry data associated with Figures 3, 5, and 6 will be deposited to the MassIVE repository and made publicly available prior to publication.

This paper does not contain original code.

Any additional information required to reanalyze the data reported in this paper is available from the lead contact upon request

### EXPERIMENTAL MODEL AND STUDY PARTICIPANT DETAILS

#### Mice

Mice were maintained in a pathogen-free facility on a 12h light, 12h dark cycle with ad libitum food and water. All procedures were reviewed and approved by the Institutional Animal Care and Use Committees (IACUC) at Yale. All breeding animals were heterozygous for the *Rbck1* null allele and had been previously backcrossed to a C57BL/6 background more than nine times. Age-matched male and female HOIL-1^+/+^ and HOIL-1^-/-^ littermates were used in all experiments.

#### Cell lines

AC16 cells were cultured in DMEM/F12, HEPES (Gibco 11320033) supplemented with 10% FBS, HEK293T (female) cells and MEFs were cultured in DMEM (Gibco 11995065) supplemented with 10% FBS, 786-O cells (male) were cultured in RPMI 1640 (Gibco 11875093) supplemented with 10% FBS, 1% penicillin-streptomycin. All cell lines were maintained at 37°C with 5% CO_2_. AC16 cells were a kind gift from Iain Scott (UPMC). HEK293T and 786-O cells were obtained from American Type Culture Collection (ATCC). HOIL-1^ΔRING1^ MEFs were previously generated (32393887). All cell lines were routinely tested for mycoplasma.

### METHOD DETAILS

#### Transverse aortic constriction

Transverse aortic constriction (TAC) was performed on mice at approximately 10 weeks of age in mice using a minimally invasive approach (26888314). After anesthesia with ketamine (100 mg/kg) and xylazine (10 mg/kg), a skin incision 0.5–1.0 cm in length was made at the level of the suprasternal notch, and a longitudinal cut 2–3 mm in length was made in the proximal portion of the sternum. This allowed for visualization of the aortic arch without opening the pleural space. A 27-gauge needle was placed next to the aortic arch to calibrate the constriction and an 8-0 suture was tied. Following ligation, the needle was removed. The skin was closed and mice were allowed to recover on a warming pad until they were fully awake.

#### Echocardiography

Mice were anesthetized with 1.5% isoflurane and 1.5% oxygen and kept on a heated platform during imaging. Echocardiography was performed with a Visual Sonics Vevo2100 ultrasound with a 40 mHz probe. Standard echo views were obtained of the left ventricle and of the aortic arch.

#### Histology

Mouse hearts were fixed in 4% paraformaldehyde for 24 hours at 4°C, then stored in 70% ethanol for an additional 48 hours. Five-micron sections were cut and parrafin-embedded. Sections were stained with hematoxylin and eosin, Masson’s Trichrome, periodic acid-Schiff, and TUNEL using standard histochemical procedure. Cardiomyocyte cross-sectional area and width were calculated using the CmyoSize macro for Fiji/ImageJ^97^. Five randomly selected regions of interest from H&E-stained heart sections at 40X magnification were used per mouse.

#### Molecular biology

All plasmid subcloning was done using the In-Fusion Snap Assembly Kit (Takara Bio 638947). Site-directed mutagenesis was done using QuikChange XL (Agilent 200517).

#### Lentivirus generation and transduction

5×10^5^ HEK293T cells were seeded on poly-L-lysine-coated 6-well plates in 2 mL DMEM+10% FBS. The following day, cells were transfected with 1.5 µg psPAX2, 0.5 µg pMD2.G, and 2.0 µg transfer vector per well using Lipofectamine 2000 according to the manufacturer’s protocol. Media was changed 8 hours after transfection. Media containing lentivirus was collected every 24 hours and stored at 4°C. After 72 hours, pooled lentiviral media was filtered through a 0.45 µm mixed cellulose esters filter (Millipore). Target cells were transduced in 6-well plates at 30-40% confluence with 1 mL lentiviral media containing 8 µg/mL polybrene. Media was replaced after 24 hours. The appropriate selection antibiotic was added to transduced cells 48 hours post-transduction. After 7-10 days of selection, transduced cells were expanded and stocks were frozen down, followed by maintenance in a half dose of the selection antibiotic. No antibiotics were included in media for experiments.

#### Knockout cell line generation

For CRISPR-Cas9 gene editing, sgRNA sequences were chosen using CHOPCHOP^98^ and cloned into either lentiCRISPRv2-puro (Addgene 52961) or lentiCRISPRv2-hygro (Addgene 98291) according to the Feng Zhang Lab protocol. Target cells were transduced as above. Following selection, cells were reseeded in 3x 96-well plates per cell line using limiting dilution. Single cell-derived colonies were expanded, and knockout efficiency was validated by immunoblot. Multiple successfully edited clones per cell line were tested in initial experiments.

#### Cell treatment and starvation

Cells were seeded such that they were at approximately 70% confluence at the time of treatment or starvation. For starvation experiments, cells were seeded on poly-L-lysine-coated plates to minimize cell loss during washes. To starve, media was aspirated and cells were washed twice quickly and gently with pre-warmed PBS, and either glucose-free DMEM (Gibco 11966025) supplemented with 10% dialyzed FBS (ThermoFisher A3382001) or Earl’s Balanced Salt Solution (Gibco 24010043) was added for glucose or amino acid starvation, respectively. For cystine starvation, media was prepared using DMEM without L-glutamine, L-cystine, glucose, phenol red, or sodium pyruvate (USBiological D9815-25L), with 4 mM L-glutamine added back fresh. L-cystine (ThermoFisher J61651.09) was added to a final concentration of 200 µM where indicated. Unless otherwise indicated, inhibitors or other supplements were added at the same time of starvation.

Transient gene silencing was achieved by reverse transfection of siRNA at a final concentration of 10 nM using RNAiMAX Lipofectamine (Invitrogen 13778075) according to the manufacturer’s protocol. All siRNA used were ON-TARGETplus SMARTpool siRNA from Horizon Discovery.

#### Glycogen quantification

9×10^4^ AC16 cells were seeded in poly-L-lysine-coated 6-well plates in triplicate per condition. The following day, cells were pre-treated with GPI (Cayman 17578) or DMSO vehicle control for 1 hour prior to glucose starvation for an additional 1 hour or left in DMEM/F12 + 10% FBS. GPI or DMSO was included during starvation. After starvation, cells were placed on ice and washed quickly with 2x 1mL ice-cold PBS, scraped into 150 µL ice-cold ddH_2_O and collected in 0.2 mL strip PCR tubes on ice. 15 µL was reserved for protein quantification and the remainder was boiled for 30 minutes at 98°C. After boiling, samples were spun down and diluted 1:10 with ddH_2_O. Glycogen was quantified using the Glycogen Assay Kit (Sigma MAK016) with slight modification: Hydrolysis Enzyme Mix was used at 1/2 the recommended concentration, Development Enzyme Mix was used at 1/3 the recommended concentration, and the Fluorescent Peroxidase Substrate was diluted 1:10. Glycogen content was normalized to protein concentration, then expressed as a percentage of glycogen measured in full media.

#### Immunoblotting

When collecting lysates for direct immunoblot analysis, media and dead cells were aspirated, and adherent cells were quickly washed with 1 mL ice-cold PBS. After removing all PBS, cells were scraped directly into modified Laemmli buffer (62.5mM Tris-HCl pH 6.8, 15% glycerol, 2% SDS, 0.025% bromophenol blue) containing 5% freshly added 2-mercaptoethanol (2-ME). Samples were boiled for 5 minutes at 95°C followed by vigorous vortexing to reduce viscosity. Samples were resolved on 4-15% or 4-20% Criterion TGX pre-cast Midi gels (Bio-Rad 5671085 or 5671095) at 90V constant voltage, followed by transfer onto 0.45 µm or 0.2 µm nitrocellulose membranes using the Trans-Blot Turbo Transfer System (Bio-Rad). Membranes were quickly washed with distilled water and blocked with 5% milk in 0.1% TBS-Tween for 1 hour at room temperature, followed by overnight incubation at 4°C in primary antibody diluted in either 5% milk in 0.1% TBS-Tween or 5% bovine serum album in 0.1% TBS-Tween. Membranes were washed 4x 10 minutes in 0.1% TBS-Tween, followed by incubation with HRP-conjugated secondary antibody in 5% milk in 0.1% TBS-Tween for 1 hour at room temperature. Membranes were washed 4x 10 minutes in 0.1% Tween and signal was detected using Amersham ECL Detection reagents (Cytiva RPN2105) or Super Signal™ West Femto (ThermoFisher 34095) with a ChemiDoc imaging system (Bio-Rad).

To examine ZAKα phosphorylation, samples were resolved on 7% acrylamide gels containing 10.7 µM Phos-Tag (Wako AAL-107) and 21.3 µM MnCl_2_ (as in Wu *et al.*^2^) and processed as above.

#### RT-qPCR

Total RNA was extracted using TRIzol reagent (ThermoFisher 15596026) according to the manufacturer’s protocol. 1 µg RNA was reverse transcribed using the High-Capacity cDNA Reverse Transcription Kit (ThermoFisher 4368814) according to the manufacturer’s protocol. cDNA was diluted 1:5 in nuclease-free water and 2 µL was mixed with 10 µL 2X SYBR Select MasterMix (ThermoFisher 4472908) and 250 nM forward and reverse qPCR primers in a 20 µL final volume per sample. Thermocycling was according to the manufacturer’s protocol. Following normalization to the housekeeping gene *TBP*, fold induction was calculated using the 2^−ΔΔCt^ method. See Table S5 for primer sequences.

#### Immunoprecipitation

For overexpression assays, 2.0×10^6^ HEK293T cells were seeded in poly-L-lysine-coated 10 cm dishes, one plate per sample. The following day, cells were transfected with 2 µg each plasmid, normalizing the total DNA payload with pCMV-Flag empty vector, using Lipofectamine 2000 at a 3:1 Lipo : DNA ratio. 48 hours post-transfection, cells were starved of glucose for 6 hours where indicated. Cells were harvested by trypsinization, centrifuged at 300 x g for 5 minutes at 4°C, and cell pellets were washed 1x 10mL ice-cold PBS. Cell pellets were lysed in 500 µL lysis buffer (10 mM Tris-HCl, 150 mM NaCl, 0.5 mM EDTA, 0.5% NP40) + fresh protease inhibitor (Thermo Scientific A32965) and left on ice for 30 minutes with intermittent gentle vortexing. Lysates were clarified at 14,000 rpm for 15 minutes at 4°C. Clarified lysates were diluted 1:2 with dilution buffer (10 mM Tris-HCl, 150 mM NaCl, 0.5 mM EDTA, protease inhibitor) and protein was quantified using the BCA protein assay (Thermo Scientific 23225). 15 µL GFP-Trap magnetic agarose beads (Proteintech gtma) were washed 3x 500 µL ice-cold dilution buffer and added to 500 µg lysate. Samples were rotated for 1 hour at 4°C, then washed 4x with wash buffer (10 mM Tris-HCl, 150 mM NaCl, 0.5 mM EDTA, 0.05% NP40). Proteins were eluted by boiling in 40 µL Laemmli containing 10% 2-ME for 10 minutes and subject to immunoblot analysis.

To examine PABPC1 ubiquitination, HEK293T cells were seeded, transfected, starved, and collected as above. Cell pellets were subjected to hot lysis by resuspending in 150 µL ubiquitin lysis buffer (1% SDS, 50 mM Tris-HCl pH 7.5, 150 mM NaCl, 5 mM EDTA) containing protease inhibitor and 50 µM PR-619 (Cayman Chemical 16276) using a wide bore pipette tip, followed by boiling for 15 minutes at 95°C with intermittent trituration. After boiling, samples were diluted 1:10 with Buffer G (1% Triton, 50 mM Tris-HCl pH 7.5, 150 mM NaCl) containing protease inhibitor and PR-619 and passed through a 22G syringe until homogenous (∼5-10 strokes). Lysates were clarified at 14,000 rpm for 15 minutes at 4°C. 500 µg clarified lysates were subjected to GFP-Trap immunoprecipitation as above.

For endogenous PABPC1 co-IP, 1.6×10^6^ AC16 cells were seeded in poly-L-lysine-coated 15 cm dishes, five plates were sample. The following day, cells were starved of glucose for 6 hours as above. Adherent cells were collected by trypsinization and centrifuged at 300 x g for 5 minutes at 4°C. Cell pellets were washed 2x 50 mL ice-cold PBS, then lysed in 1 mL lysis buffer (10 mM Tris-HCl, 150 mM NaCl, 1.0 mM EDTA, 0.5% NP40 + fresh protease inhibitor) and left on ice for 30 minutes with intermittent gentle vortexing. The same 1 mL was used to harvest all five plates of a sample. Lysates were clarified at 14,000 rpm for 15 minutes at 4°C. For each sample, 4 µg anti-PABPC1 (Proteintech 10970-1-AP) or rabbit IgG isotype control (Cell Signaling Technology 3900S) was added to 50 µL washed Dynabeads Protein G beads (Invitrogen 10003D) in 1.5 mL LoBind tubes (Eppendorf 022431081) containing 200 µL PBS + 0.02% Tween, mixed well, and rotated at room temperature for 40 minutes. Antibody-bead complexes were washed 1x 200 µL 0.02% PBS-Tween, then 2 mg lysate in 1 mL final volume was added and rotated overnight at 4°C. Beads were washed 4x 200 µL ice-cold 0.02% PBS-Tween and transferred to a clean tube with the last wash. Proteins were eluted by boiling in 40 µL Laemmli containing 10% 2-ME for 10 minutes and subject to immunoblot analysis.

#### RNA granule fractionation

Fractionation was performed as in Namkoong *et al*.^79^. 9×10^5^ (for 24h glucose starve) or 4.5×10^5^ (for untreated, 8h glucose starve, and sodium arsenite treatment) AC16 cells were seeded in poly-L-lysine coated 15-cm dishes, three plates per sample. 24h glucose starvation was initiated when cells reached 80% confluence. The following day, cells were starved of glucose for 8h, treated with 0.5 mM sodium arsenite for 1h in complete media, or left untreated. After treatment, media was aspirated and cells were washed 2x 10 mL ice-cold PBS, then scraped into 1 mL ice-cold Buffer L (50 mM Tris pH 7.5, 50 mM NaCl, 5 mM MgCl_2_, 0.1% NP-40, fresh 1 mM 2-ME, 100 U/mL RNase inhibitor (Sigma 3335402001), protease and phosphatase inhibitors) and mechanically lysed by 30 strokes with a Dounce homogenizer on ice. The same 1 mL lysis buffer was used for all plates of a sample. 50 µL was removed and mixed with 50 µL Laemmli buffer + 10% 2-ME as the whole cell extract fraction. The remaining 950 µL was centrifuged at 2,000 x g at 4 °C for 2 minutes. The resulting pellet (nuclear fraction) was washed 1x 300 µL complete Buffer L then resuspended in 300-400 µL Laemmli + 5% 2-ME, while the supernatant was centrifuged again at 10,000 x g at 4°C for 10 minutes. The resulting pellet (RNA granule fraction) was washed 1x 300 µL complete Buffer L, then resuspended in 40-60 µL Laemmli + 5% 2-ME. The supernatant (cytosolic fraction) was mixed with 1:1 with Laemmli + 10% 2-ME. All samples were frozen at − 20°C until immunoblot analysis.

#### Cell death analysis

To study cell death, cells were seeded in 12-well plates such that they were at 70% confluence at the time of starvation or treatment the following day (6×10^4^ AC16 cells, 7×10^4^ 786-O cells, 3×10^4^ MEFs). After the indicated treatment, cells were harvested by collecting the media containing floating cells in 5 mL round bottom polystyrene tubes (Corning 352052). Adherent cells were washed once with room temperature PBS, trypsinized, and pooled with the floating cells. Cells were centrifuged at 2,400 rpm for 5 minutes at 4°C. The supernatant was removed and cell pellets were resuspended in 250 µL PBS containing 5 µg/mL propidium iodide (ThermoFisher P3566) and stained for 15 minutes at room temperature in the dark. Samples were processed on a BD LSRFortessa X-20 Cell Analyzer. Data were analyzed using FlowJo.

#### LC-MS/MS global metabolomics

AC16 cells were seeded in Primaria 6-well plates (Corning 353846) such that they were at 70% confluence at time of starvation (1×10^5^ cells/well for 24 h starvation, 6×10^4^ cells/well for 3 h starvation) in sextuplicate in 2 mL DMEM/F12 supplemented with 10% FBS. Cells were starved by carefully aspirating media, washing cells twice with 1 mL room temperature D-PBS, then adding 2 mL pre-warmed DMEM without glucose, glutamine, or phenol red (ThermoFisher A1443001) supplemented with 10% dialyzed FBS and 4 mM L-glutamine. After starvation, plates were removed from the incubator one plate at a time and placed on ice. Media was aspirated and cells were washed quickly with 1 mL ice-cold 5 mM HEPES. 150 µL ice-cold quenching buffer (20% methanol into 0.1% formic acid, 3mM sodium fluoride, 5.5 µg/mL D8-Phenylalanine) was added to each well. Cells were quickly scraped into quenching buffer and transferred to a pre-chilled V-bottom polypropylene plate (BD 53263) on dry ice. Once all samples were collected, the plate was sealed with aluminum film (Bioexpress T-3025-8B) and stored at −80°C.

Cell lysates were lyophilized in the 96-well plate. Samples were prepared by resuspending the cell pellet in 50 µL 10% acetonitrile solution with D_4_-taurine (25 µM) as a second internal standard. 5 µL of the supernatant was injected for each analysis mode on the mass spectrometer.

##### Chromatography

Two columns were used separately: Thermo Scientific Hypercarb column (100 x 4.6 mm, 3 µm) and a Phenomenex Kinetex F5 Core-shell LC column (100 x 2.1 mm, 2.6 µm).

Hypercarb column (HC column): Separations were performed with 1 mL/min linear gradients as indicated (Table 1). Mobile phase A: 15 mM ammonium formate, 0.03% acetyl acetone and 0.1% formic acid; mobile phase B: 60% ACN, 35% IPA, 15 mM ammonium formate and 0.1% formic acid. Column temperature was 50 °C and autosampler temperature was 5 °C.

**Table 1:**
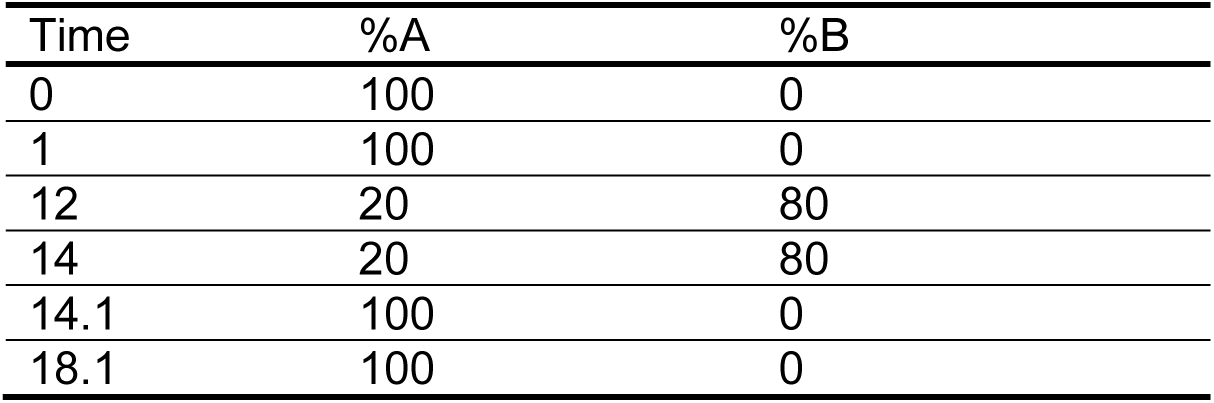
Hypercarb column chromatographic separation.

Kinetex F5 column (RP column): Separations were performed with 0.3 mL/min linear gradients as outlined (Table 2). Mobile phase A: 95% water, 5% acetonitrile and 0.1% formic acid; mobile phase B: 95% acetonitrile, 5% water and 0.1% formic acid. Column temperature was 30 °C and autosampler temperature was 5 °C.

**Table 2:** Kinetex F5 column chromatographic separation.

| Time | %A | %B |
| --- | --- | --- |
| 0 | 100 | 0 |
| 0.5 | 100 | 0 |
| 15 | 0 | 100 |
| 16 | 0 | 100 |
| 16.1 | 100 | 0 |
| 20 | 100 | 0 |

##### Mass spectrometry

Data were analyzed using the Sciex TripleTOF 6600 collected using an information-dependent analysis (IDA) workflow consisting of a TOF MS scan (200 msec) and a high-resolution IDA experiment (70 msec each) monitoring 10 candidate ions per cycle. Former target ions were excluded after 2 occurrences for 5 sec and dynamic background subtraction was employed. The mass range for both TOF MS and IDA MS/MS scans was 60-1000 with RP column and 70-1000 with HC column.

The ion source conditions were as follows; Ion spray voltage = 5000 V for positive mode coupling with RP column and −4500 for negative mode coupling with HC column, ion source gas 1 (GS1) = 50, ion source gas 2 (GS2) = 50, curtain gas (CUR) = 30, temperature (TEM) = 400 °C with RP column and 500°C with HC column. Compound dependent parameters for the two modes were: declustering potential (DP) = 35, collision energy (CE) = 30, collision energy spread (CES) = 20.

##### Data processing

El-MAVEN software (Elucidata.io) was used for peak picking and curation from house built targeted and untargeted libraries. Targeted libraries used the commercial standard kit of ∼ 600 metabolites (IROA Technologies) in the 2 modes of analyses (RP column positive ion acquisition mode, HC column negative ion acquisition mode) yielding reference data (molecular ions and retention times) with wide coverage of the endogenous metabolome. Untargeted libraries contain ∼2,700 metabolites drawn KEGG database, and the top 5 candidates with the highest intensity throughout a sample run for each metabolite were automatically curated.

Data were normalized by the sum of all targeted metabolites within each sample in each mode and then log transformed. Sample quality was assessed using two internal standards, D_8_-phenylalanine and D_4_-taurine to identify any outliers to be excluded from the downstream analysis. Next, the normalized data from the two modes were merged. If a metabolite appeared in both modes, the one with the best intensity was chosen.

PCA, PLS-DA and heat map from the merged targeted data were performed using MetaboAnalyst (https://www.metaboanalyst.ca/). Differential expression analysis for each comparison was performed using the metabolomics application in Polly (polly.elucidata.io). Normal p values and log2FC data was uploaded in Shiny GAM (https://artyomovlab.wustl.edu/shiny/gam/) to generate an optimized metabolic network displaying ∼60 of the most changing and closely connected metabolites. The up-regulated and down-regulated metabolites in the treatment groups were then used separately for pathway enrichment analysis in MetaboAnalyst (https://www.metaboanalyst.ca).

#### NADP+/NADPH quantification

4×10^3^ AC16 cells were seeded in poly-L-lysine-coated 96-well plates. The following day, cells (at ∼60% confluence) were starved of glucose by aspirating media, washing wells twice with 50 µL room temperature PBS, then gently adding 75 µL glucose-free DMEM supplemented with 10% dialyzed FBS. 16 mM glucose was added to the media when indicated. NADP/NADPH levels were measured using the NADP/NADPH-Glo Assay (Promega G9081) following the manufacturer’s protocol.

#### di-GG TMT-based proteomics and ubiquitinomics

1.8×10^6^ AC16 cells were seeded in poly-L-lysine-coated 15 cm dishes, three plates per sample in quadruplicate. When cells reached 80% confluence, they were starved of glucose as above. After starvation, cells were washed 2x 10 mL ice-cold PBS and scraped into 1 mL urea lysis buffer (9 M urea, 50 mM HEPES pH 8.5, 0.5% sodium deoxycholate) with 20 mM 2-chloroacetamide (Sigma C0267) and 4 µg/mL Lys-C (Wako 12505061). The same 1 mL was used to harvest all three plates of a sample. Samples were immediately flash frozen in liquid nitrogen then stored at −80°C

For protein extraction, samples were gently thawed, then placed in a Bullet Blender at a speed of 6 for 30 s with an interval of 10 s at 40C for 6 cycles. Proteins were digested with Lys-C (200:1 protein : enzyme ratio by weight, Wako 12505061) in the presence of 10 mM DTT and 5% acetonitrile for 3 h, followed by a 4-fold dilution with 50 mM HEPES buffer pH 8.5 and overnight trypsin digestion (50:1 protein : enzyme by weight, Promega V5280) at room temperature. The peptides were desalted on C18 SPE column (Waters) and dried under vacuum.

The enrichment of di-Gly peptides was performed using PTMScan® HS Ubiquitin/SUMO Remnant Motif (K-GG) Kit (Cell Signaling Technology) as previously described^99^. Briefly, the di-Gly peptides were isolated from 2 mg peptides by magnetic K-GG beads after incubation at 40°C for 1.5 h. The eluted di-Gly peptides were adjusted to pH 8.5 using 50 mM HEPES and labeled with 60 μg TMTpro reagent at room temperature for 30 mins. After the reaction was quenched with 5% hydroxylamine, the TMTpro labeled di-Gly peptides were mixed in equal amounts, desalted on C18 stagetips and fractionated in a 60 min gradient of 15-50% B on a reverse phase HPLC column (ACQUITY UPLC BEH C18 column, 1.0 x 100 mm, 1,7 μm particle size, Waters) on an Agilent 1220 HPLC system (buffer A: 10 mM ammonium formate, pH 8.0; buffer B: 90% MeCN, 10 mM ammonium formate, pH 8.0) and collected into ten fractions. All fractions were dried, reconstituted in 5% formic acid and analyzed using acidic pH reverse phase LC-MS/MS (CoAnn Technologies, 75 μm ID x 20 cm, 1.9 μm C18 resin at 60°C) on an Ultimate 3000 UPLC system (Thermo Fisher Scientific). Peptides were eluted in a 120 min gradient of 16%-55% buffer B (67% MeCN, 3% DMSO, and 0.1% FA) with buffer A (3% DMSO, and 0.1% FA) at a flow rate of 0.25 µL /min before MS analysis.

For total proteome analysis, the peptides in the flowthrough from di-Gly enrichment experiments were labeled with TMT pro at a ratio of 1.5:1 by weight. After mixing in equal amounts, the fractionation was performed on an ACQUITY UPLC BEH C18 column (2.1 x 100 mm, 1.7 μm particle size, Waters). The HPLC gradient was from 15-50% B for 120 min at a flow rate of 0.2 ml/min on the same Agilent 1220 HPLC system. A total of 40 pooled fractions were dried under vacuum, resuspended in 5% formic acid, and analyzed by acidic pH reverse phase LC-MS/MS in an 80 min gradient of 16%-55% buffer B before MS analysis.

The Orbitrap HF mass spectrometer (ThermoFisher Scientific) was operated in data-dependent mode. The parameters for MS1 scan are set as 60,000 resolution, scan range 460–1600 m/z, 3 × 106 AGC target, and 50 ms maximal ion time for whole proteome analysis. The parameters for MS2 scans are 120 m/z low mass threshold, 1 × 105 AGC, 110 ms maximal ion time, 20 data-dependent MS2 scans, 1.0 m/z isolation window with 0.2 m/z offset, 32 normalized collision energy in HCD, and 10 s dynamic exclusion for whole proteome analysis. For ubiquitinome analysis, MS1 scans are set as 60,000 resolution, scan range 530–1,600 m/z, 3 × 10^6^ AGC target, and 50 ms maximal ion time. MS2 scans are set as 120 m/z low cutoff, 1 × 10^5^ AGC, 150 ms maximal ion time, 17 data-dependent MS2 scans, 1.0 m/z isolation window with 0.2 m/z offset, 30-32 normalized collision energy in HCD, and 20 s dynamic exclusion.

The mass spectrometric database search was processed by the JUMP software suite^100^. The MS/MS spectra were searched against the UniProt human protein database, which contained 170,122 entries and was downloaded in 2020. The search parameters included a mass tolerance of 15 ppm for both precursor ions and MS/MS ions, full trypticity with a maximum two missed cleavages, a maximum of three dynamic modification sites per peptide, dynamic modification with Met oxidation (+15.99492 Da) and Lys di-Gly modification (+114.04293 Da, for ubiquitinome), and static modifications with TMT tags on Lys and N-termini (+304.2071453 Da), and Cys carbamidomethylation (+57.02146 Da). All protein and peptide identifications were filtered based upon mass accuracy and matching scores using the target-decoy strategy to reduce protein or peptide FDR to < 1%. The quantification of proteins and di-Gly peptides was performed using JUMP suite pipeline. Differentially expressed di-Gly peptides were determined using both p values and log2 fold change (FC) calculated by the limma R package. The significance of differential expression was evaluated based upon statistical criteria, with p < 0.01 and Log2(FC) > two-fold of the standard deviation (SD). The SD of di-Gly peptides was estimated by fitting to a Gaussian distribution to evaluate the magnitude of experimental variations.

#### Polysome profiling

1.8×10^6^ AC16 cells were seeded in poly-L-lysine-coated 15 cm dishes, four plates per sample. Cells were starved of glucose for 8 hours as above or treated with 5 µM anisomycin for 15 minutes in full media. After treatment, cells were washed 3x 10 mL ice-cold PBS + 100 µg/mL cycloheximide (CHX) on ice. Cells were scraped into 1 mL polysome lysis buffer (15 mM Tris-HCl pH 7.5, 5 mM MgCl_2_, 150 mM NaCl, 1% Triton X-100) supplemented with fresh 2 mM DTT, 100 µg/mL CHX, and protease inhibitor. The same 1 mL was used to harvest all plates of a sample. Lysates were left on ice for 5 minutes, then clarified at 14,000 rpm for 5 minutes at 4°C. All samples were diluted to 1.7 mg/mL in 1 mL final volume and flash frozen in liquid nitrogen.

10-50% sucrose gradients (15 mM Tris pH 7.5, 5 mM MgCl_2_, 150 mM NaCl, 2 mM DTT, 100 µg/ml CHX) were made and fractionated using Biocomp Gradient Station (Biocomp Instruments). Lysates were thawed on ice, added to sucrose gradients and centrifuged at 36,000 RPM at 4°C in a Beckmann SW41 rotor for 2 hours. During fractionation, absorbance was continually monitored at 260 nm. Fractions were collected and subjected to immunoblot analysis.

#### Proximity labeling

1.8×10^6^ MiniTurbo fusion-expressing AC16 cells were seeded in poly-L-lysine-coated 15 cm dishes, two plates per sample. When cells reached ∼70% confluence, media was aspirated, cells were washed 2x 10 mL PBS and replaced with 20 mL glucose-free DMEM + 10% dialyzed FBS + 50 µM D-biotin (Sigma-Aldrich B4639). 16 mM D-glucose was added to control plates. After starvation, cells were washed 2x 10 mL ice-cold PBS, then scraped into 1 mL RIPA lysis buffer (150 mM NaCl, 5 mM EDTA pH 8.0, 50 mM Tris-HCl pH 8.0, 1% NP-40, 0.5% sodium deoxycholate, 0.1% SDS) containing protease and phosphatase (Sigma-Aldrich 4906845001) inhibitors. The same 1 mL was used to harvest all plates of a sample. Lysates were placed on ice for 30 minutes with intermittent vortexing, then clarified at 14,000 for 20 minutes at 4°C. Clarified lysates were pre-cleared with 15 µL washed Protein A magnetic beads (Cell Signaling Technology 73778) for 30 minutes at 4 °C with rotation. 50 µL washed Streptavidin Magnetic beads (Pierce 88816) were added to 1.5 mg lysate in 1 mL final volume and rotated overnight at 4°C. The next day, beads were washed 4x 1mL ice-cold RIPA, followed by 4x 1mL ice-cold PBS. The beads were transferred to a fresh tube on the last wash. PBS was completely removed and the beads were subjected to LC-MS/MS analysis.

### QUANTIFICATION AND STATISTICAL ANALYSIS

Statistical analyses were performed using GraphPad Prism software (Version 10). Bar graphs are the mean ± standard error from one representative experiment. The number of replicates (n) used in an experiment is indicated in the figure legend. Two-tailed Student’s t-test was used for evaluating statistical significance between two groups. Two-way analysis of variance (ANOVA) followed by Sidak’s post-hoc comparison was used to evaluate statistical significance between two or more independent parameters. Asterisks denote significance: *p<0.05, **p<0.01, ***p<0.001, ****p<0.0001, ns: p>0.05

## SUPPLEMENTAL INFORMATION

See Figures S1-S8 and Tables S1-S5 (Excel files).

## SUPPLEMENTAL FIGURE LEGENDS

**Figure S1.**
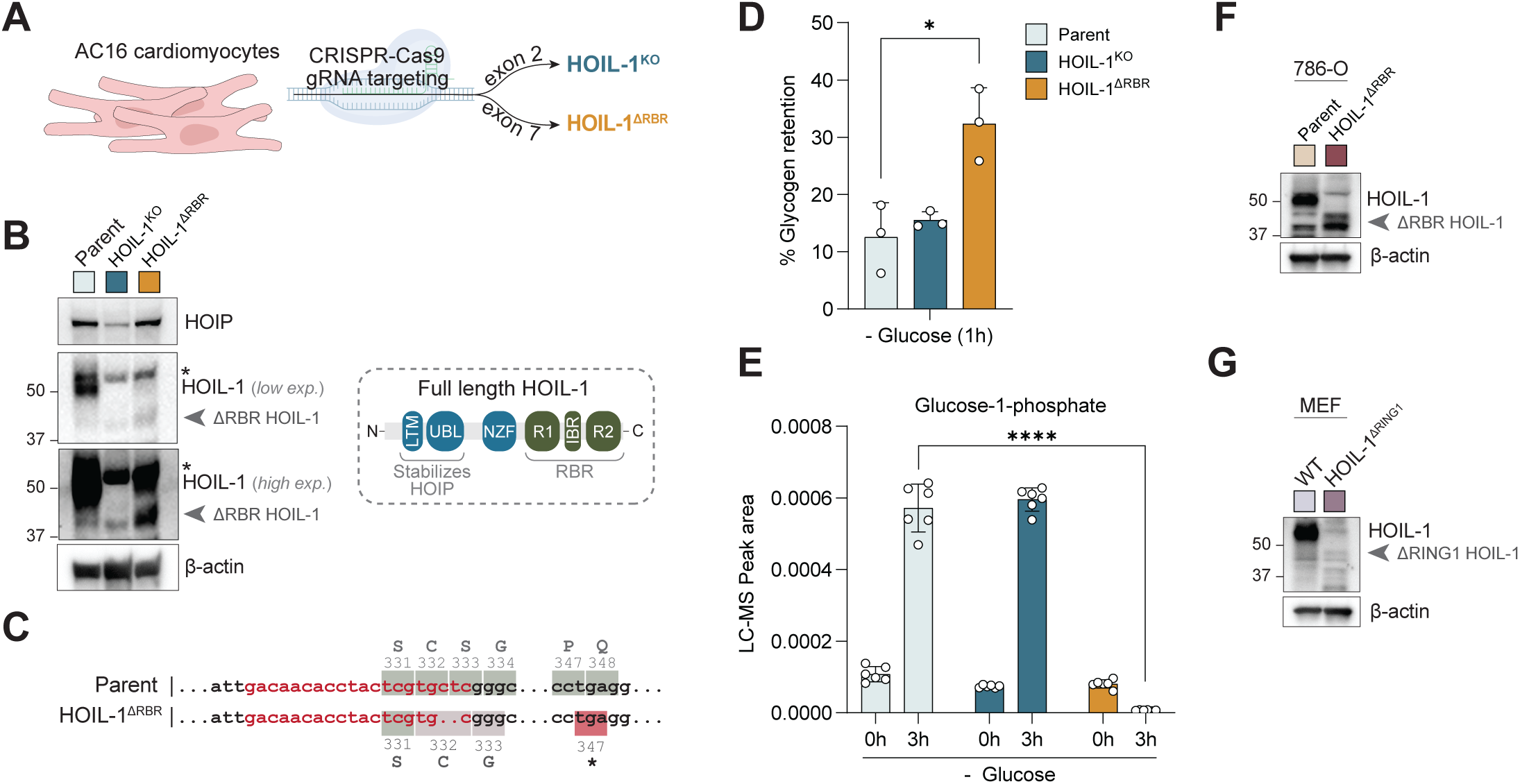
Loss of the HOIL-1 RBR impairs glycogen breakdown. (A) Model system. (B) Immunoblot analysis of AC16 cells. Arrowhead indicates ΔRBR HOIL-1. Asterisks indicate non-specific bands. (C) Genomic DNA sequencing from AC16 cells confirming deletion and premature stop codon in HOIL-1^ΔRBR^ cells. Red text is the gRNA target sequence. (D) Fraction of cellular glycogen remaining after 1h glucose starvation in AC16 cells. Pooled data from 3 experiments. Each data point is the mean of 3 replicates (n=3). (E) Glucose-1-phosphate levels in AC16 cells after glucose starvation measured by LC-MS (n=6). (F) Immunoblot analysis of HOIL-1^ΔRBR^ 786-O cells. (G) Immunoblot analysis of HOIL-1^ΔRING1^ MEFs.

**Figure S2.**
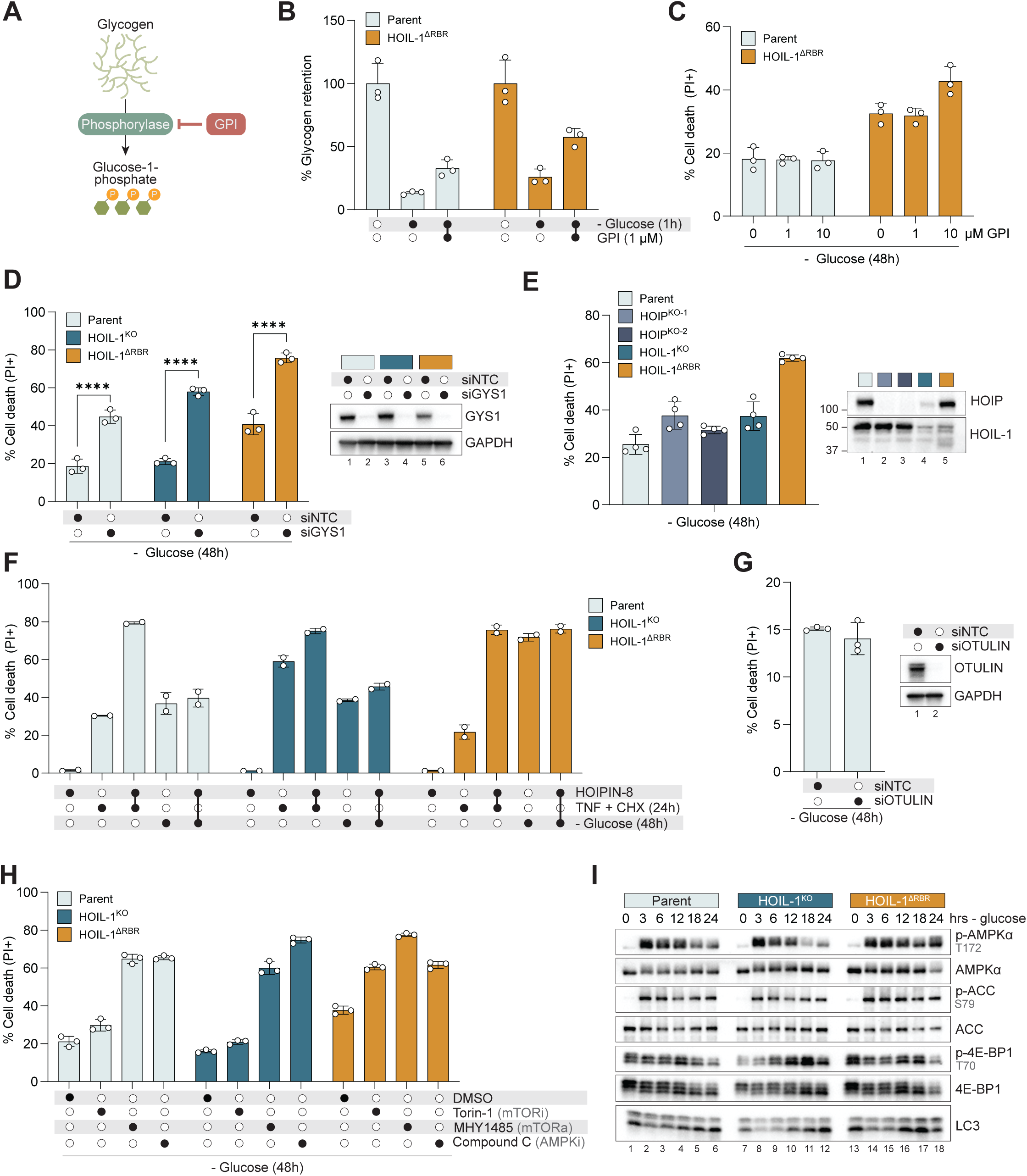
Glucose starvation causes noncanonical cell death. (A) Schematic. Glycogen phosphorylase inhibitor (GPI) reduces glycogen breakdown. (B) Fraction of cellular glycogen remaining after 1h glucose starvation in AC16 cells pre-treated with GPI (1 µM) for 1h. (C) Cell death in AC16 cells pre-treated with GPI for 1h followed by 48h glucose starvation (n=3). (D) Cell death in AC16 cells transfected with GYS1 or non-targeting control (NTC) siRNA 48h prior to glucose starvation for 48h (n=3). (E) Cell death in HOIP^KO^ AC16 cells after 48h glucose starvation (n=4). (F) Cell death in AC16 cells TNF (20 ng/mL) + CHX (6 µg/mL) treatment for 24h or glucose starvation for 48h in combination with HOIPIN-8 (30µM) (n=2). (G) Cell death in AC16 cells transfected with OTULIN or NTC siRNA 48h prior to glucose starvation for 48h (n=3). (H) Cell death in AC16 cells treated with DMSO vehicle, Torin-1 (250 nM), MHY1485 (20 µM), or Compound C (7 µM) concurrent with 48h glucose starvation (n=3). (I) Immunoblot analysis for AMPK and mTOR-related proteins in AC16 cells following glucose starvation for the indicated time-points.

**Figure S3.**
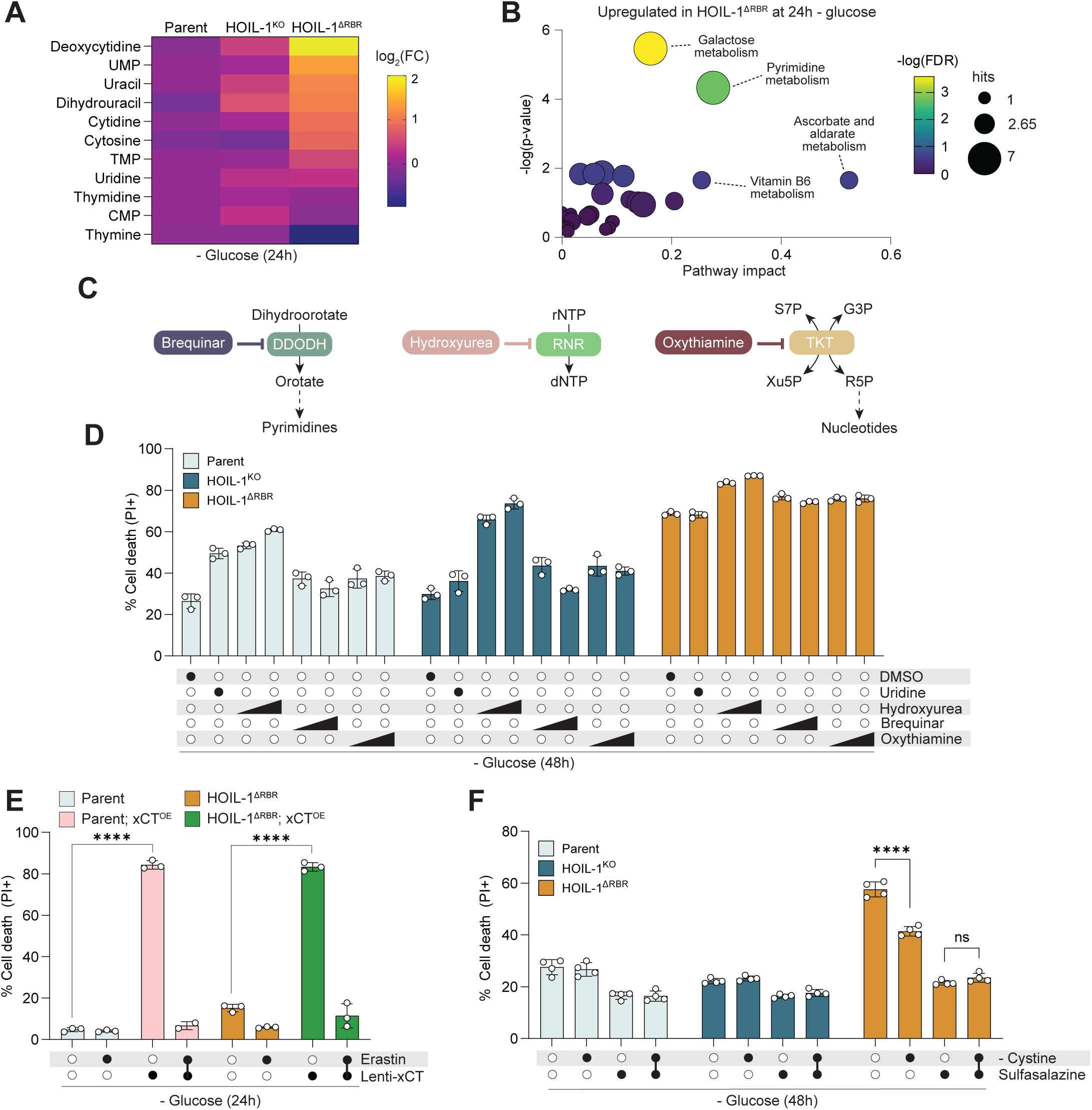
Pyrimidines do not enhance glucose starvation-induced cell death. (A) Heatmap of metabolite levels after 24h represented as log2(FC) of parental AC16 cells. (B) Pathway analysis of metabolites enriched in HOIL-1^ΔRBR^ cells after 24h glucose starvation. (C) Schematic of inhibitors of pyrimidine metabolism. (D) Cell death in AC16 cells after DMSO vehicle, uridine (25 mM), hydroxyurea (0.2 mM, 2.0mM), brequinar (0.1 µM, 1.0 µM), or oxythiamine (0.5 mM, 5 mM) treatment concurrent with 48h glucose starvation (n=3). (E) Cell death in AC16 cells stably overexpressing (OE) xCT treated with erastin (2 µM) concurrent with 48h glucose starvation (n=3). (F) Cell death in AC16 grown in L-cystine and glucose-free media supplemented with L-cystine (200 µM) or sulfasalazine (500 µM) for 48h (n=4).

**Figure S4.**
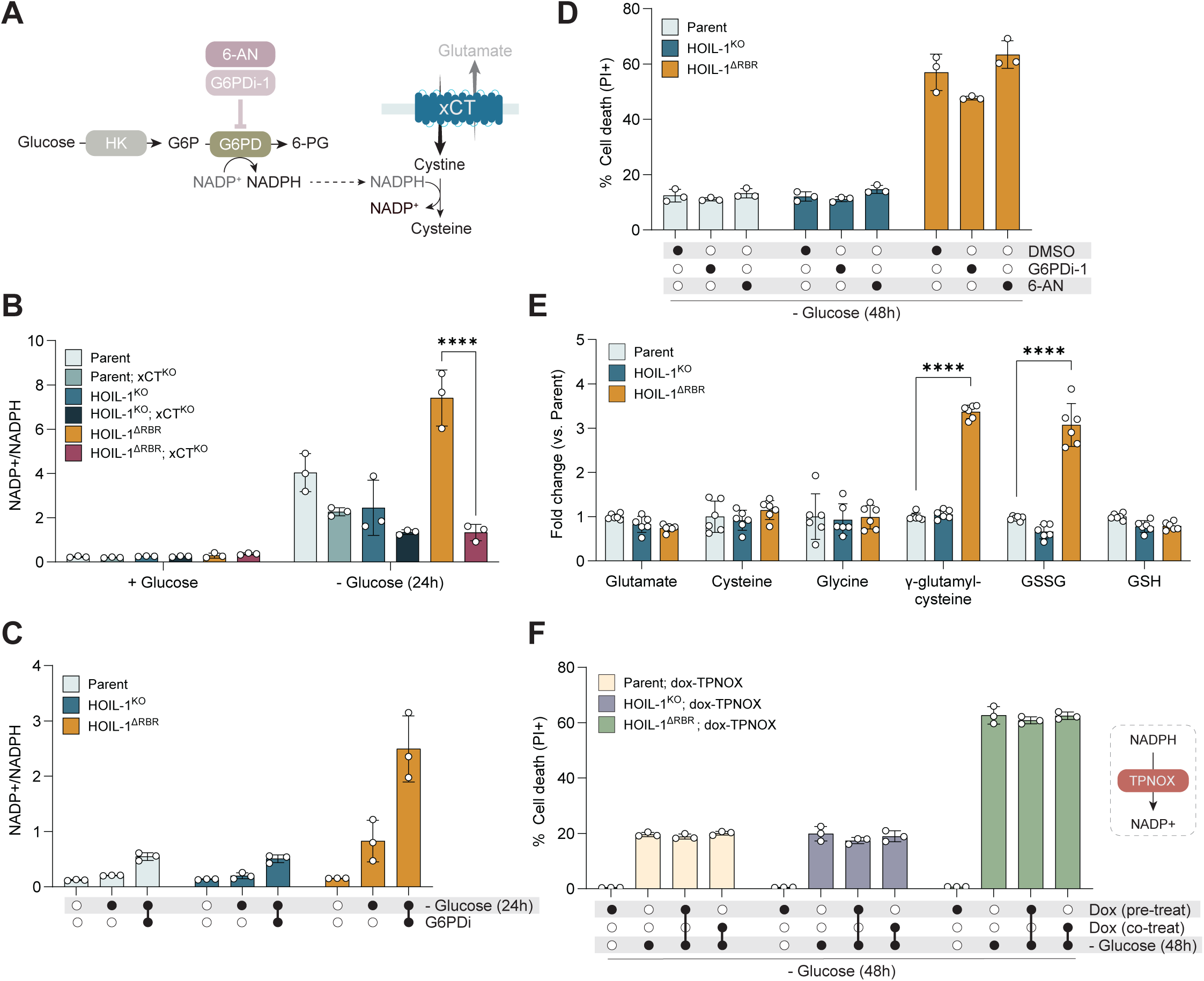
NADPH collapse does not cause glucose starvation-induced cell death. (A) Schematic of NADP+/NADPH flux. (B) Measurement of NADP+ and NADPH ratio during glucose starvation in xCT^KO^ AC16 cells (n=3). (C) Measurement of NADP+ and NADPH ratio in AC16 cells treated with G6PD inhibitor (G6PDi-1, 50 µM) during glucose starvation (n=3). (D) Cell death in AC16 cells after DMSO vehicle, G6PDi-1 (50 µM), or 6-aminonicotinamide (6-AN, 20 µM) treatment concurrent with 48h glucose starvation (n=3). (E) Cell death in AC16 cells stably overexpressing doxycycline (dox)-inducible TPNOX treated with dox (1 µg/mL) for either 24h prior to (pre-treat) or concurrently (co-treat) with 48h glucose starvation (n=3). (F) xCT-related metabolite levels at 24h glucose starvation as measured by untargeted LC-MS represented as fold change over parent (n=6).

**Figure S5.**
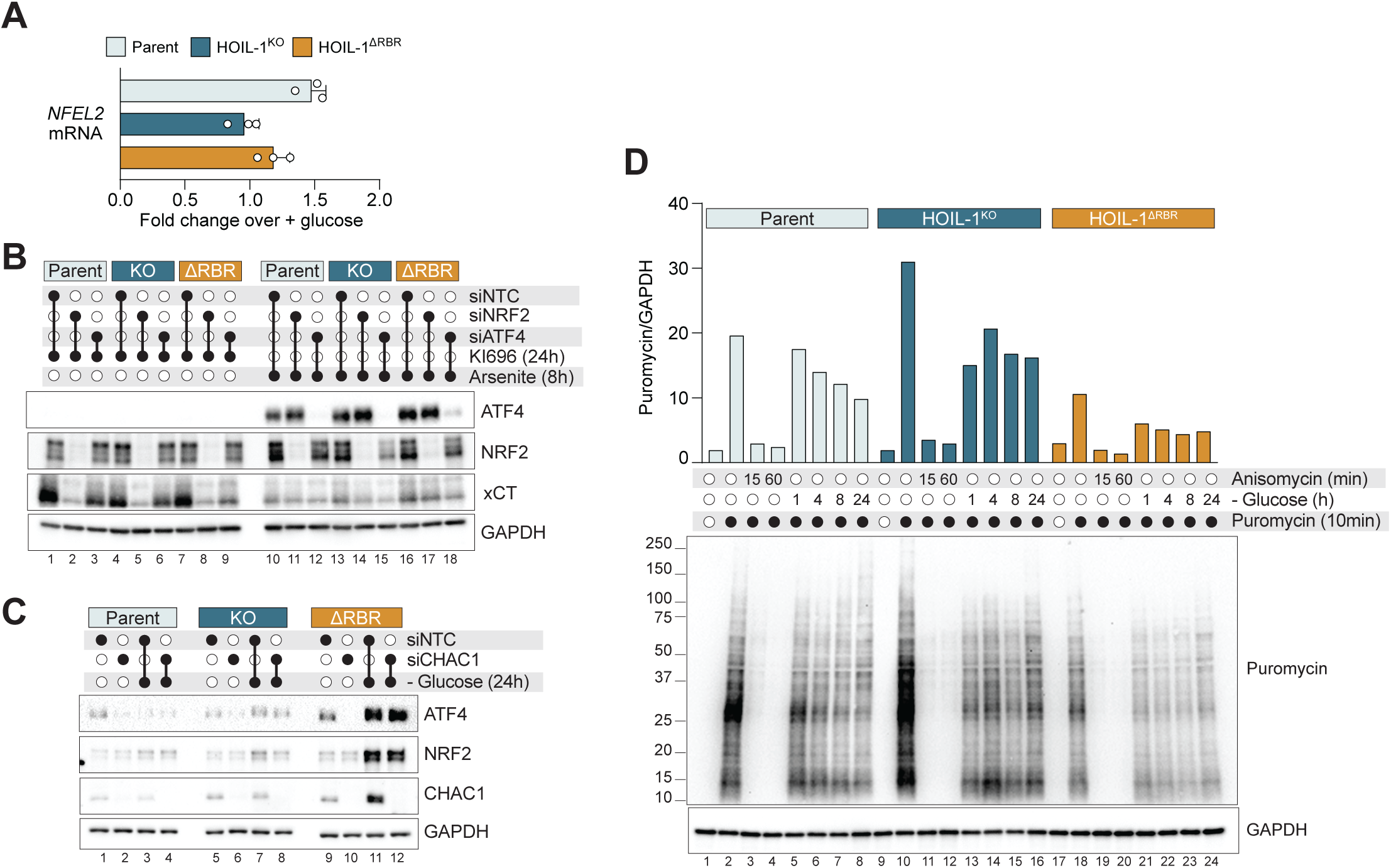
ATF4 post-transcriptionally regulates NRF2. (A) NRF2 (*NFEL2)* mRNA levels (normalized to *TBP*) in AC16 cells after 24h glucose starvation. Data represented as the fold change over full media (n=3). (B) Immunoblot analysis of AC16 cells transfected with ATF4, NRF2, or non-targeting control (NTC) siRNA 48h prior to treatment with KI696 (NRF2 activator, 1µM) for 24h or sodium arsenite (10µM) for 8h. (C) Immunoblot analysis of AC16 cells transfected with CHAC1 or non-targeting control (NTC) siRNA 48h prior to glucose starvation for 24h. (D) AC16 cells were treated with anisomycin (5µM) or starved of glucose for the indicated times. 10 minutes prior to lysis, cells were treated with 10µg/mL puromycin. Immunoblot analysis was performed using an anti-puromycin antibody and densitometric analysis was performed on puromycin and GAPDH signals.

**Figure S6.**
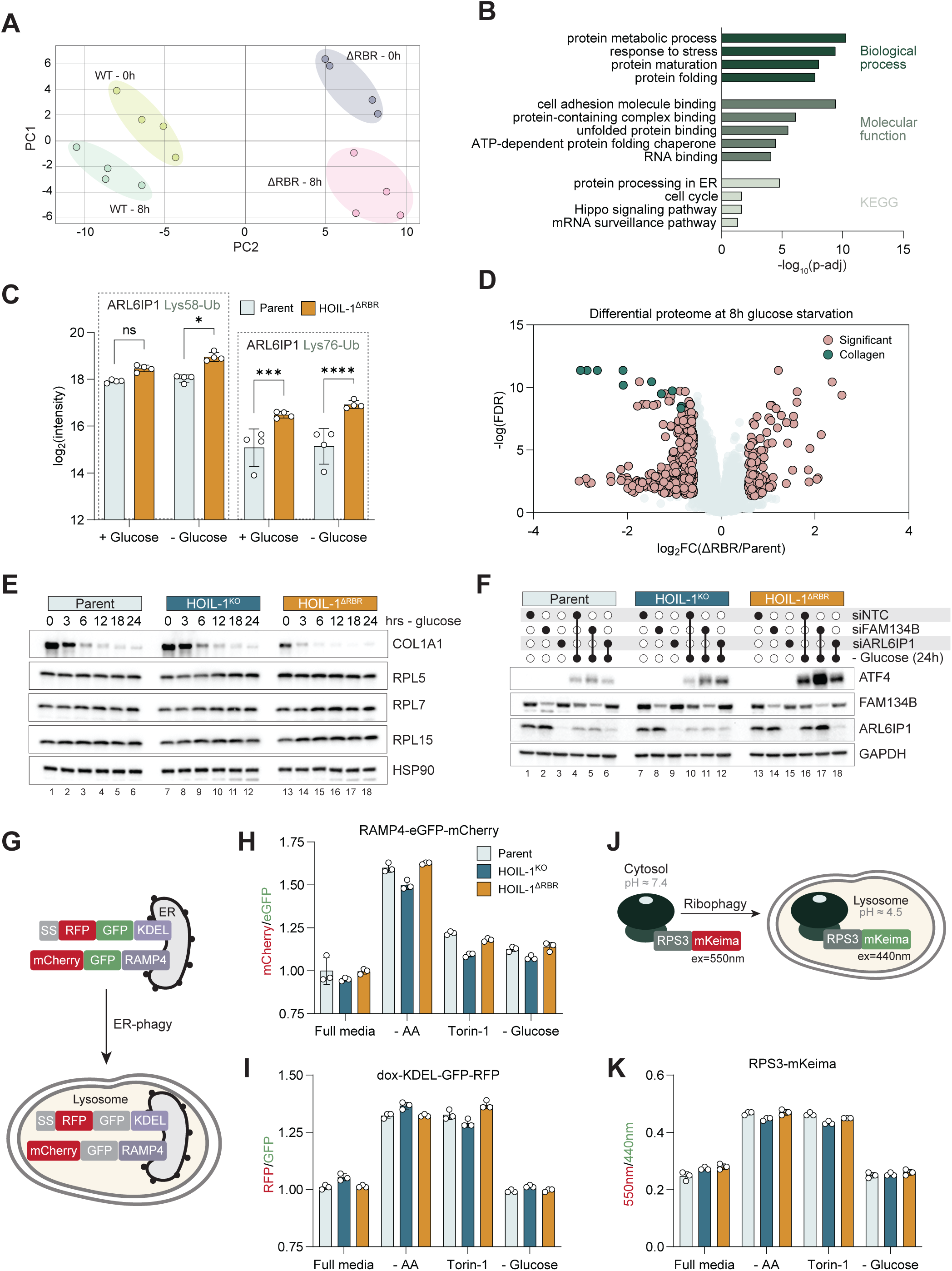
Selective autophagy is not involved in the glucose starvation response. (A) Principal component analysis of ubiquitinome data. (B) Gene ontology enrichment analysis of proteins with increased ubiquitin-modified peptides in HOIL-1^ΔRBR^ cells relative to parent AC16 cells after 8h glucose starvation. (C) Log transformed intensities of ARL6IP1 ubiquitinated lysines in AC16 cells detected by LC-MS/MS (n=6). (D) Volcano plot of differentially abundant proteins in parent and HOIL-1^ΔRBR^ AC16 cells after 8h glucose starvation. Significant: FDR > 0.05, log2FC > 2SD. (E) Immunoblot analysis of AC16 cells starved of glucose for the indicated time-points. (F) Immunoblot analysis of AC16 cells transfected with FAM134B, ARL6IP1, or non-targeting control (NTC) siRNA 48h prior to glucose starvation for 48h. (G) Schematic of ER-phagy reporters. GFP quenching in the acidic lysosome alters the GFP:RFP ratio. (H) Flow cytometric analysis of mCherry and GFP intensity in AC16 cells stably expressing RAMP4-eGFP-mCherry treated with Torin-1 (250nM) or starved of amino acids (AA) or glucose for 24h (n=3). (I) As in (G) but cells stably expressing dox-KDEL-GFP-RFP were treated with doxycycline (1 µg/ml) for 24 hours prior to starvation or treatment (n=3). (J) Schematic of ribophagy reporters. (K) As in (G) and (H) but cells are expressing RPS3-mKeima (n=3).

**Figure S7.**
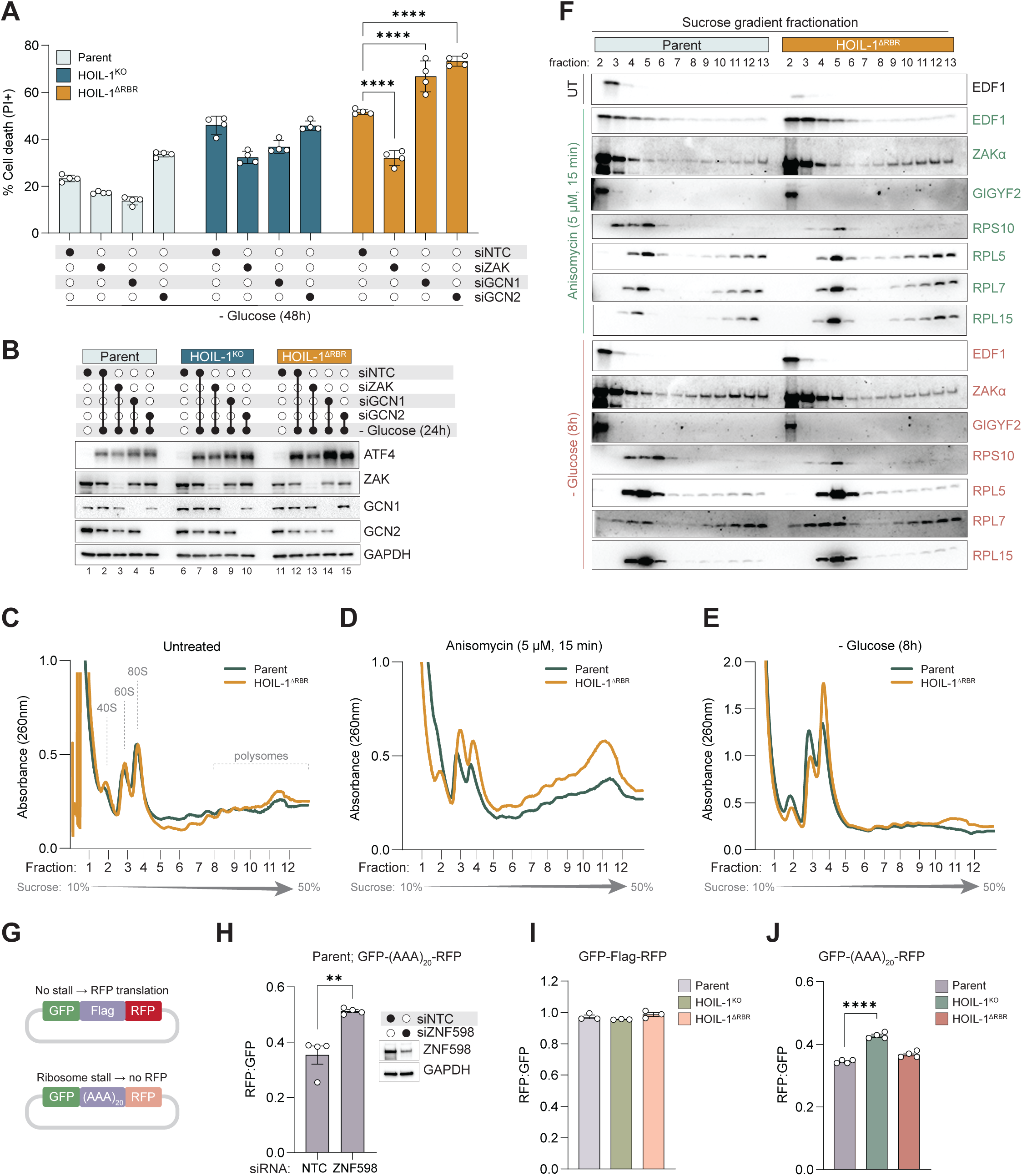
Glucose starvation activates ZAKα in HOIL-1^ΔRBR^ cells through a noncanonical mechanism. (A) Cell death in AC16 cells transfected with ZAK, GCN1, GCN2 or non-targeting control (NTC) siRNA 48h prior to glucose starvation for 48h (n=4). (B) Immunoblot analysis of AC16 cells transfected with ZAK, GCN1, GCN2 or NTC siRNA 48h prior to glucose starvation for 24h (C) Polysome profiles from lysates of untreated AC16 cells (D) Polysome profiles from lysates of AC16 cells treated with anisomycin (5µM) for 15 minutes (E) Polysome profiles from lysates of AC16 cells starved of glucose for 24h. (F) Immunoblot analysis of sucrose fractions from (C)-(E). (G) Schematic of ribosome stalling reporter constructs. Ribosome stalling on (AAA)_20_ reduces RFP translation. (H) Flow cytometric analysis of GFP and RFP in AC16 cells stably expressing the ribosome stalling reporter transfected with ZNF598 or NTC siRNA 48h prior (n=4). Immunoblot analysis of ZNF598 knockdown shown. (I) Flow cytometric analysis of GFP and RFP in AC16 cells stably expressing the ribosome nonstalling reporter (n=3). (J) Flow cytometric analysis of GFP and RFP in AC16 cells stably expressing the ribosome stalling reporter (n=3).

**Figure S8.**
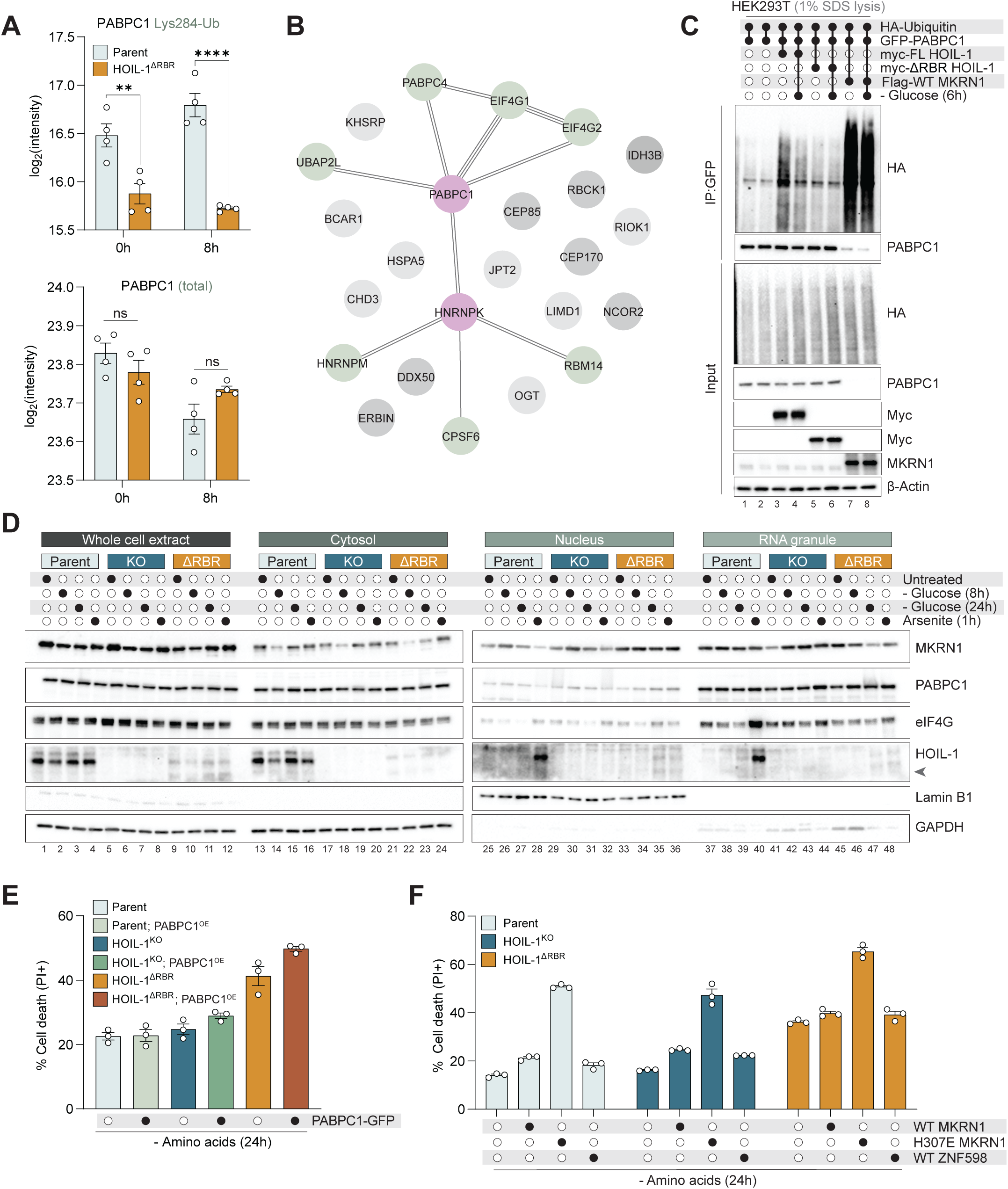
HOIL-1 ubiquitinates PABPC1 and controls MKRN1 localization. (A) Log transformed intensities of PABPC1 ubiquitinated Lys284 and total levels in AC16 cells detected by LC-MS/MS (n=6). (B) STRING analysis of the ΔRBR HOIL-1 proximitome during glucose starvation. (C) Immunoblot analysis of HEK293T cells transfected with the indicated plasmids 24h prior to 6h glucose starvation where indicated, followed by hot lysis in 1% SDS and anti-GFP immunoprecipitation. (D) Immunoblot analysis of AC16 cells treated with sodium arsenite (10µM) or starved of glucose for the indicated time-points, followed by subcellular fractionation of RNA-protein granules. Arrowhead indicates ΔRBR HOIL-1. (E) Cell death in AC16 cells stably expressing PABPC1-GFP after 24h amino acid starvation (n=3). (F) Cell death in AC16 cells stably expressing MKRN1 or ZNF598 after 24h amino acid starvation (n=3).

